# Anticipatory responses along motion trajectories in awake monkey area V1

**DOI:** 10.1101/2020.03.26.010017

**Authors:** Giacomo Benvenuti, Sandrine Chemla, Arjan Boonman, Laurent Perrinet, Guillaume S Masson, Frédéric Chavane

## Abstract

What are the neural mechanisms underlying motion integration of translating objects? Visual motion integration is generally conceived of as a feedforward, hierarchical, information processing. However, feedforward models fail to account for many contextual effects revealed using natural moving stimuli. In particular, a translating object evokes a sequence of transient feedforward responses in the primary visual cortex but also propagations of activity through horizontal and feedback pathways. We investigated how these pathways shape the representation of a translating bar in monkey V1. We show that, for long trajectories, spiking activity builds-up hundreds of milliseconds before the bar enters the neurons’ receptive fields. Using VSDI and LFP recordings guided by a phenomenological model of propagation dynamics, we demonstrate that this anticipatory response arises from the interplay between horizontal and feedback networks driving V1 neurons well ahead of their feedforward inputs. This mechanism could subtend several perceptual contextual effects observed with translating objects.

**Highlights:**

- Our hypothesis is that lateral propagation of activity in V1 contributes to the integration of translating stimuli
- Consistent with this hypothesis, we find that a translating bar induces anticipatory spiking activity in V1 neurons.
- A V1 model describes how this anticipation can arise from inter and intra-cortical lateral propagation of activity.
- The dynamic of VSDi and LFP signals in V1 is consistent with the predictions made by the model.
- The intra-cortical origin is further confirmed by the fact that a bar moving from the ipsilateral hemifield does not evoke anticipation.
- Horizontal and feedback input are not only modulatory but can also drive spiking responses in specific contexts.

## INTRODUCTION

In standard information processing models of the visual system, low-level visual features are extracted locally within selective receptive fields and are rapidly relayed to downstream areas in order to encode increasingly complex features (Fukushima, 1980; Riesenhuber and Poggio, 1999). When probed with spatially stationary stimuli centered on their receptive fields, properties of visual neurons from V1 to extra-striate areas are indeed well captured by a simple cascade of *linear-nonlinear models* encoding low-level visual properties such as local orientation or motion direction (Carandini et al., 2005; Rust et al., 2006) as well as more complex visual pattern (David et al., 2006; DiCarlo et al., 2012). A well-known example is given by *visual motion integration* where three decades of studies have been conducted almost exclusively with motion signals presented within a stationary aperture, such as drifting periodic stimuli or random dot patterns, centered on neuronal receptive fields (**RF**). The reported neuronal properties have provided strong evidences for the *linear-nonlinear cascade* theoretical framework, where V1 neurons encode local ambiguous motion signals that are integrated by MT neurons to compute direction and speed of pattern motion (Movshon et al., 1985; Rust et al., 2006; Simoncelli and Heeger, 1998; Wilson et al., 1992).

However, natural objects often move along extended and smooth spatio-temporal trajectories within the visual field. A handful of psychophysical, computational and physiological studies (Watamaniuk & Mc Kee 1995, Watamaniuk et al 1995, Burgi et al., 2000; Perrinet and Masson, 2012; Tlapale et al., 2010) have proposed that the visual system can use the spatio-temporal regularity of smooth trajectories to solve many of the uncertainties inherent to visual motion such as aperture (Marr, 1982; Wallach, 1935) and correspondence problems (Ullman, 1979; Wertheimer, 1912) and to improve the *signal to noise ratio* of visual input evoked by moving stimuli (e.g. Burgi et al., 2000). This idea is consistent with the view that visual systems can benefit from contextual information and internal models of the statistics of the visual world (Berkes et al., 2011; Geisler, 2008).

For example, the property of smoothness associated to motion trajectories can be used by the visual system to generate internal models of moving objects (i.e. priors). In accordance with this hypothesis, there is a rich psychophysical literature demonstrating how spatio-temporal coherence affects the way we perceive moving stimuli attributes such as (1) position (MacKay, 1958; Nijhawan, 1994; W. Metzger, 1932), (2) direction (Anstis and Ramachandran, 1987; Lorenceau et al., 1993; Ramachandran and Anstis, 1983; Welch et al., 1997), (3) motion (Van Doorn and Koenderink, 1984; Nakayama and Silverman, 1984; Neri, 2014; Snowden and Braddick, 1989b; Verghese and Watamaniuk, 1999; Verghese et al., 2000; Watamaniuk and McKee, 1995; Watamaniuk et al., 1995), (4) speed (Castet et al., 1993; Georges et al., 2002; McKee and Welch, 1985; Scott-Brown and Heeley, 2001) (5) orientation (Alais and Lorenceau, 2002; Fredericksen et al., 1994; Guo et al., 2004; Pavan et al., 2011; Werkhoven et al., 1990) and (6) color (Watanabe and Nishida, 2007). For example, a target stimulus presented at the leading edge of a smooth trajectory has been shown to be easier to detect than when presented in isolation or at the trailing edge of the trajectory (Arnold et al., 2014; Lenkic and Enns, 2013; Liu et al., 2006; Roach et al., 2011; Schwiedrzik et al., 2007). This phenomenon cannot be simply explained by a shift in spatial attention, lateral facilitation or retinal motion and it is more likely implemented at an early stage of visual processing (Roach et al., 2011). Furthermore, stimuli presented at the leading edge of long smooth trajectories are perceived having higher resolution (i.e. lower blurring due to motion) than stationary moving stimuli (Burr and Ross, 1986; Watanabe and Nishida, 2007) and are more easily trackable behind occlusions (Watamaniuk and McKee, 1995).

Several computational studies suggest that a simple hierarchical feedforward model with contiguous local filters sequentially activated by the translating stimulus is insufficient to account for these contextual effects. In contrast, they proposed that a *mechanism involving lateral propagation across the sequence of filters activated along the trajectory* could explain the emergence of motion trajectory effects (Burgi et al., 2000; Grzywacz et al., 1995; Kaplan et al., 2013; Khoei et al., 2017; Perrinet and Masson, 2012; Tlapale et al., 2010; Yuille and Grzywacz, 1989). These models postulate that lateral interactions in early visual areas could participate in processing the responses evoked by smooth trajectories of translating stimuli. Early visual areas are “retinotopically organized” (Van Essen and Newsome, 1984; Hubel and Wiesel, 1974), such that translating stimuli generate a sequential activation of contiguous locations on the retinotopic map. Since neurons in these areas are tightly connected to their neighbors through horizontal connections and feedback loops (Angelucci et al., 2002; Benucci et al., 2007; Bringuier et al., 1999; Bullier, 2001; Chavane et al., 2011; Girard et al., 2001; Grinvald et al., 1994; Jancke et al., 2004a; Muller et al., 2014; Reynaud et al., 2012; Slovin et al., 2002) see review (Muller et al., 2018) (**Figure 1A**), feedforward responses at each location can rapidly spread to contiguous areas of the map before the translating stimulus enters their RF (Jancke et al 2004, Chemla et al 2019). Consequently, the neural population mapping the leading edge of the translating stimulus will receive convergent lateral and FF inputs. These two signals may be recursively integrated along the motion trajectory and generate a gradual build-up of neural activity that would carry trajectory information. The advantage of such integration mechanism is that it would combine only the spatio-temporally coherent responses evoked by translating stimuli.

**Figure 1.**
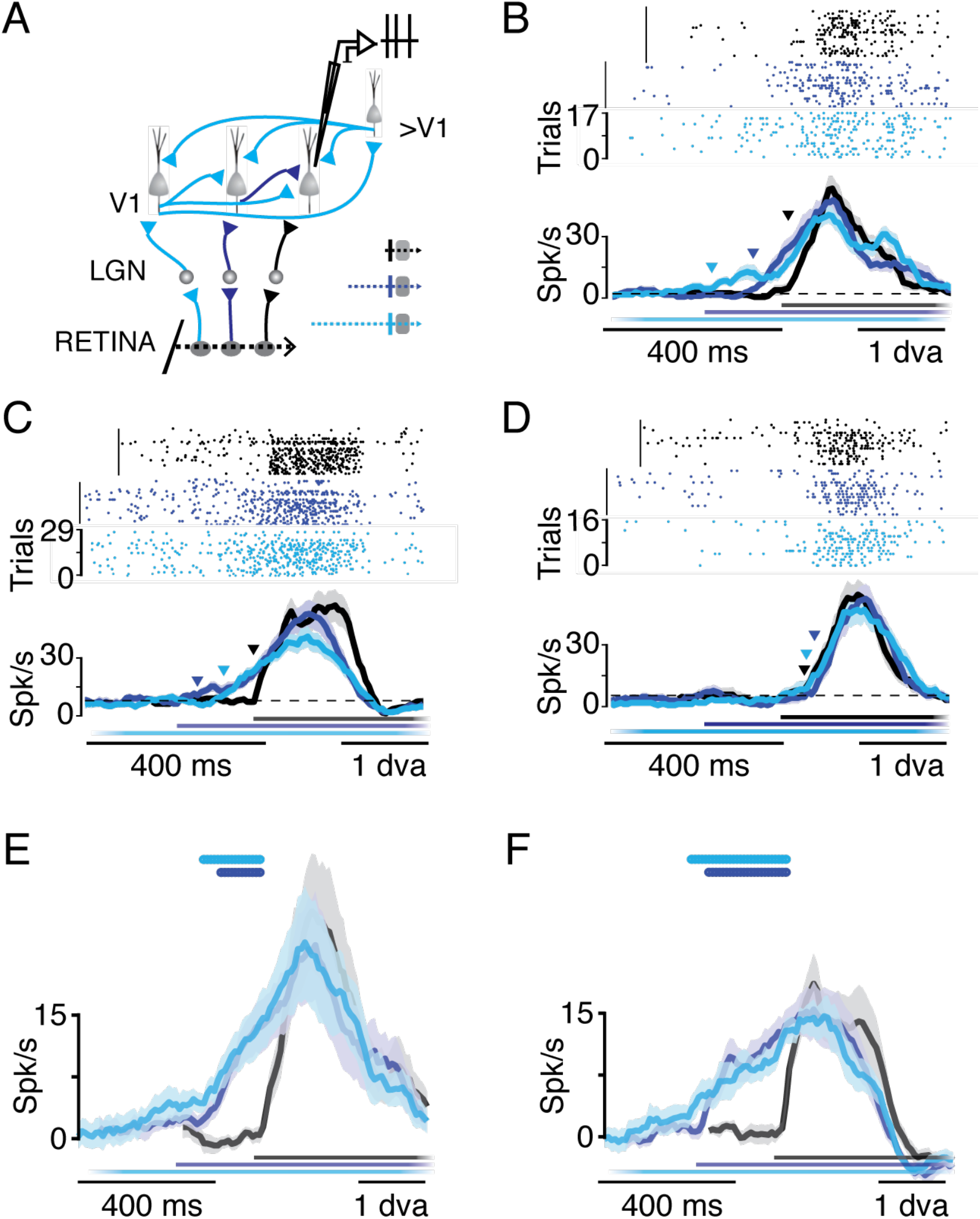
Single units responses to a translating bar as a function of its trajectory length outside the RF. (**A**) Experimental approach. Schematic of the retino-thalamo-cortical neural circuit. A translating bar (bottom) is going to cross consecutive V1 RFs (grey disks) eliciting a sequence of feedforward activations, propagating laterally across the retinotopic map through intra (i.e. horizontal) and inter-cortical (feedback-loops) connections. Neurons with the RF at the end of the trajectory are recorded extracellularly using an electrode (top-right). The inset on the right depicts the paths of the three trajectories (dotted lines, 1.5, 3 and 6dva, respectively) with respect to the RF (grey circle). The colorcode is consistent throughout figures. (**B-D**) Examples of single unit responses for the three trajectory lengths presented with raster plots (top) and PSTHs (bottom) (B and D: Monkey PD, C: Monkey NO). B and C are examples of neurons presenting anticipation, D of a neuron that does not. Colored lines (bottom) indicate the stimulus presentation time, for each condition. Latency is indicated by triangles. Shades represent the standard error of the mean (SEM) across trials. (**E-F**) Average PSTHs across anticipatory neurons for the three trajectories, for monkeys PD (n=13) and NO (n=13), respectively. Responses are aligned to the latency of the short trajectory condition. Colored lines (top) indicate significant difference with respect to the short trajectory condition (two-sample t-test P<0.05).

The aim of this study is to investigate this intriguing hypothesis. The primary visual cortex (**V1**) is a good candidate circuit to support such trajectory-based computation as it has the largest and most precise retinotopic representation of the outside world (Hubel and Wiesel, 1974; Tootell et al., 1982). To understand how V1 networks map motion trajectories, it is thus necessary to elucidate the interactions between feedforward, feedback and lateral networks at the proper spatio-temporal recording scales.

To achieve this goal, we recorded both single-unit and neuronal population activities in the primary visual cortex of four awake, fixating, monkeys to map the dynamics of the cortical responses to a translating bar. By using different trajectory lengths, we manipulated the history of cortical activation elicited by the moving bar before reaching the classical RF of the recorded neuron. Using single-unit recordings, we found that for long trajectories, about half of V1 neurons demonstrate a significant build-up of anticipatory spiking responses long before the bar enters their classical receptive field. Using a computational model, we show that the specific dynamics of spiking anticipatory responses could be explained by an intra-cortical horizontal and feedback network propagating information faster than the feedforward input sequence. Using voltage-sensitive dye imaging (**VSDi**) to measure the aggregate membrane potential of a large V1 area, we showed indeed that the neural population response to a translating bar exhibits a spatio-temporal structure consistent with the predictions of the model. To test whether feedback signals could also contribute to this anticipatory activity, we used a multi-electrode array to measure local field potential (**LFP**) responses across the V1 area and showed that a bar moving over a long trajectory elicits a strong and very early decrease of low frequency power, a signal recently attributed to feedback modulation (Bastos et al., 2014; Engel and Fries, 2010; van Kerkoerle et al., 2014). The intra-cortical origin of these anticipatory responses was further demonstrated by the observation that anticipation was abolished when the bar translation was initiated from the ipsi-lateral visual hemi-field. Altogether, these results dissect out the multiple scales at which an intricated network of connectivities within and between visual areas leads to the emergence of an anticipatory spread of activity that shall contribute to processing of translating objects.

## RESULTS

The goals of this study were to test whether trajectory-dependent responses to a translating bar can be observed in V1 and, if so, understand the underlying network mechanisms.

### Anticipatory spiking responses

We first recorded spiking responses from 80 V1 single-units in two alert, fixating macaques presented with three visual stimulation paradigms. Neurons were selected regardless of their tuning properties. For each neuron, we first mapped the spatial profile of the classical RF using sparse noise sequences and the orientation and direction tunings, using a translating bar (12 directions). Then, a vertical bright bar (0.5×4 degrees of visual angle, **dva**) was moved horizontally at 6.6 dva/s starting at either 1.5, 3 or 6dva (short, medium and long trajectories respectively) from the RF centre and disappearing 1.5-2dva beyond it (**Figure 1A**). Critically, these three trajectories stimulated the RF in exactly the same way. However, in the medium and long trajectory conditions, the translating bar had a different history, activating a large area outside of the RF through a smooth path before reaching it.

For each cell, the responses for the three trajectory conditions were aligned relative to the absolute position of the bar (**Figure 1B-F**). On the other hand, for comparing the evoked responses across cells (**Figure 1E-F**), we aligned all time-courses relative to the latency of the response evoked by the short path condition. This operation led to the automatic exclusion of 31 cells for which the detection of this important reference point was not possible due to low signal-to-noise ratio (see Methods) and the fact that stimulus direction and orientation were not optimized for the cells’ preference.

In the remaining 49 cells, the time-course of the responses (PSTH) to the shortest (1.5dva) motion trajectory presented a typical fast transient rise and a slower decrease (**Figure 1B-F**, black trace). However, when the bar trajectory started 3dva or 6dva before the RF centre (dark and light blue curves, respectively), a ramp-like increase of the discharge-rate was observed before the response-onset for the shortest motion trajectory, in half (53%) of the neurons (**Figure 2A**). Such anticipatory build-up scaled up with the trajectory length. In the other neurons, responses were identical for the three trajectory conditions (**Figure 1D & figure supplement 1**), excluding that anticipatory responses were due to a systematic bias in our specific experimental approach (e.g. stimulation protocol, general bias due to eye movements, generalized allocation of spatial attention, generalized effect due to the input from LGN). On average, latencies for the medium (3dva) and long (6dva) trajectories started respectively 116ms (std =93ms) and 220ms (std = 189ms) before the latency to the shortest path (1.5dva), a decrease which was found significant over the whole population (two-sample t-test P<0.05, see **Figure 2D**). This timing corresponded to an average distance of 2.3 and 3dva between the bar position and the RF centre. In contrast, both time-to-peak and response offset remained similar across all trajectories (two-sample t-test P>0.05), excluding a spatial shift of the RF or an unspecific gain effect.

**Figure 2.**
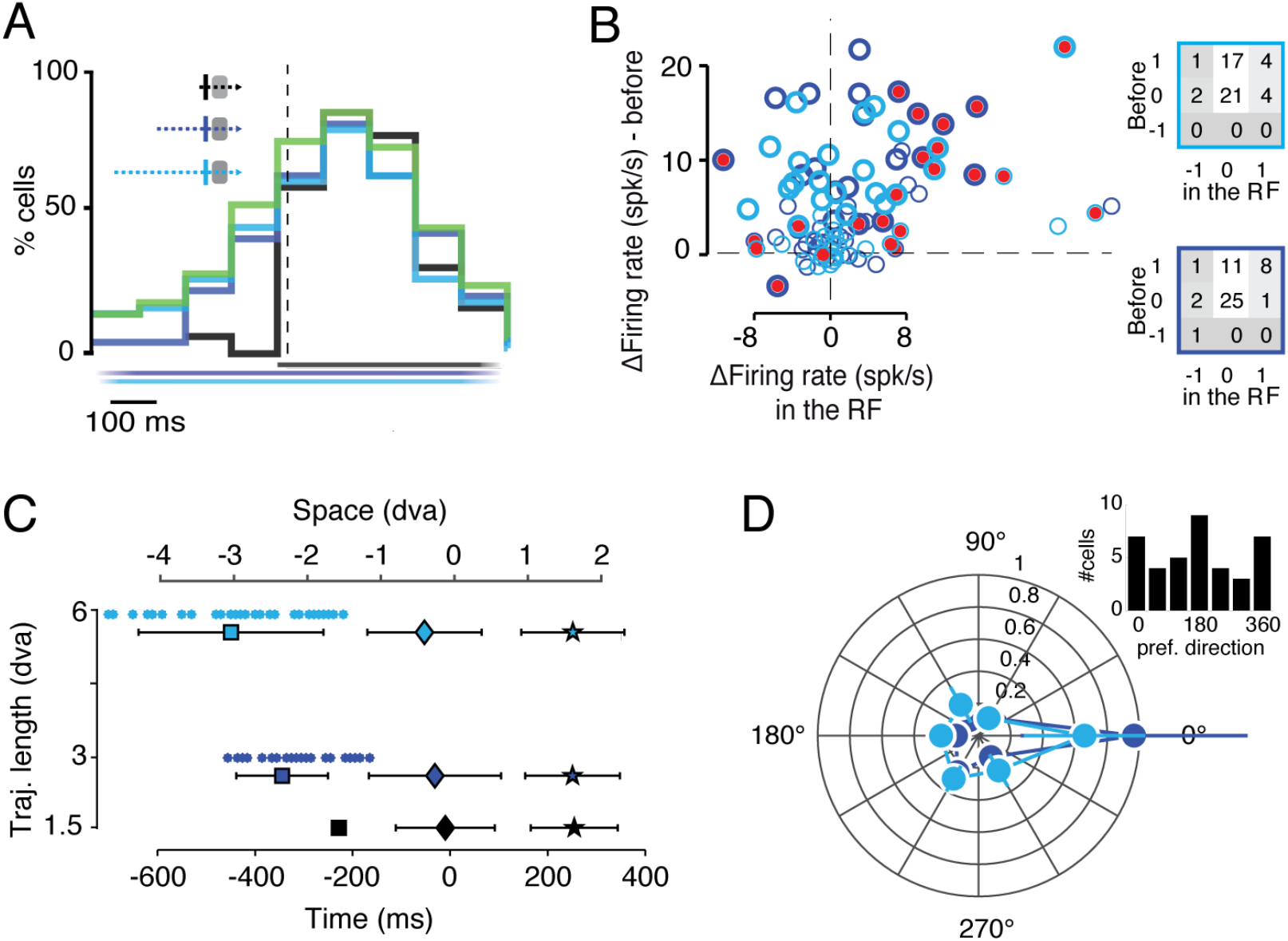
Amplitudes, latencies and tunings of trajectory dependent responses. (**A**) Proportion of neurons (out of a total of n=49) showing a significant response over time for all 3 trajectory conditions (see cartoon in insert). Green color represents proportion of neurons showing a significant response only for the medium and/or long trajectory conditions. Colored lines at the bottom indicate stimulus presentation for the three trajectories. (Top-left) same inset than **Fig1A**. (**B**) Difference of firing rate between responses to long (3-6dva) and short (1.5dva) trajectory lengths before entering the RF (thick when significant) vs within the RF (red filled when significant). Before and within the RF was estimated using the latency of the short trajectory response. Table on the top: number of cells presenting a significant increase (1), decrease (−1) or no change (0) in firing-rate rate in long trajectories compared to short trajectories. Table on the bottom: same than table on the top but for medium trajectories (**C**) Mean onset latencies (square), peak response times (diamond) and offset response time (stars) for the three trajectories conditions as a function of trajectory length averaged across neurons and monkeys (n=49). Time 0 is when the bar crosses the RF center. Dots indicate latencies of individual medium and long trajectories responses. An second abscissae is drawn at the top to indicate the equivalent visual distance (dva) from the receptive field center. (**D**) Polar histogram representing the normalized averaged anticipatory response strength (computed in a time window 100ms before the latency of the short trajectory) as a function of neurons direction preference (n=32, only cells with DI>0.1). Inset: number of cells preferring each direction.

This anticipatory activity extended well beyond the size of the measured RF, similar to classical results (Angelucci et al., 2002), and was neither correlated with the RF sizes nor could be explained by a systematic bias in monkeys’ fixation (see **figure supplement 2**). The gradual rise of this anticipatory response, furthermore, contrasts with the transient profile of the neural response for a bar swept over the RF and suggests a slower cumulative process. These anticipatory responses merged ~100ms before the short-trajectory response onset and from then followed an identical dynamic. This asymmetry eliminates the possibility that the response buildup would be related to a systematic misestimation of the classical RF position or size, that would have resulted in a change in response peak time for instance. Next, we aimed at better characterizing the origin of the cell-to-cell variability in the observed anticipatory responses, and its relationship to the tuning of the cells.

### Understanding the diversity of degrees of anticipation

To estimate the proportion of cells showing a significant modulation of firing rate over time, we compared spike counts across trials in short (i.e. 1.5 dva) *vs* long (3 & 6 dva) trajectories conditions. The proportion of cells significantly activated (Two-sample t-test P<0.05) at different time bins along the motion trajectory gradually increased (**Figure 2A**), such that 53% of them showed a significant anticipatory response 100ms before the bar entered the classical RF, for long (3dva and/or 6dva, green) but not short (1.5dva, black) trajectories. By comparison, when the bar entered the classical RF (i.e. vertical dotted line), all three conditions drove the same number of cells.

In a few neurons, the two longest trajectories resulted in a significantly different spiking response in the RF center (e.g. the example in **Figure 1C**). This result could provide an important evidence in support of the predictive coding theory, as prediction would result in smaller error signal and hence a smaller neuronal response-(Alink et al., 2010; Mumford, 1992; Srinivasan et al., 1982). However, this effect was not evidenced in the average response profiles (**Figure 1E-F**) although it could have been smoothed out because of the variability in response latencies. Hence, we compared individual discharge rates between short and long trajectories in two different time-windows of 100ms: one placed just before (ordinate) and one just after (abscissae) response latency (as estimated by detecting the first significant change with respect to the baseline) for the short trajectory condition (i.e. the RF border) (**Figure 2B**). A small fraction of the cells exhibiting an anticipatory build-up of activity responded slightly less when the bar crossed their RF center (upper left quadrant in Figure 2B). However, such decrease was significant in only 4% of the cells while, on the contrary, 16% displayed a significant increase of their response in the RF (insets). These results rule out a systematic relationship between the occurrence of an anticipatory build-up and a modulation of the stimulus-driven responses within the RF center.

Next, we compared, for every neuron, the latencies for medium and long trajectories vs. the latency for the short one (**Figure 2C**). This analysis revealed a continuum in the timing of anticipation as shown by the latency scatter for the medium and long trajectories that fully covered a wide range of values from 0 to −400 and −800ms, respectively. While the distributions of response onset latencies were significantly different for the three trajectories conditions (two-sample t-test P<0.05) both time-to-peak and response offset latencies distributions were not significantly different across all trajectories (two-sample t-test p>0.05). We checked if the variability in anticipatory responses across all neurons can be due to differences in their tuning properties. Selecting neurons with a direction index > 0.1 we probed the relationship between the relative firing rate in the 100ms time-window before the bar enters the RF and direction preferences (**Figure 2D**). Indeed, neurons with a preferred direction aligned to the stimulus motion direction exhibited a stronger anticipatory response. Therefore, the diversity of the anticipation timing and strength of the response across neurons may simply reflect a direction-selective anticipatory mechanism.

Overall, we demonstrate that a large population of V1 neurons exhibits a slow build-up of activity prior a moving input hits their receptive field. Such anticipatory activity occurs sooner for longer trajectories (**Figure 2C,** squares). In comparison, both the response peak and offset response time did not change with trajectory length (Figure 2C, diamond and stars respectively). Our next goal was to understand what neural mechanisms could generate this anticipatory response.

### Anticipation is an expected emergent property of intra-cortical propagation

These characteristics of observed anticipatory responses can be used to theoretically constrain the putative underlying neural mechanisms. First, these responses emerge well before the stimulus enters the classical RF of the recorded neurons. Second, the decrease in onset latency is not followed by a similar shift in time-to-peak and offset latency. Eventually, anticipatory responses present a slow dynamic consisting of a gradual rise of activity. A feedforward network alone is unlikely to account for the slow dynamics and the asymmetry of these responses. Alternatively, these features are consistent with surround modulation (Bair et al., 2003; Klink et al., 2017; Nassi et al., 2013; Reynaud et al., 2012). Visual information from the RF surround can be provided to V1 neurons from intra-cortical horizontal (**H**) and inter-cortical feedback (**FB**) networks (Angelucci and Bressloff, 2006; Roelfsema, 2006). We therefore reasoned that the anticipation we observed in V1 shall arise from the interactions between feedforward and intra-cortical inputs carried by horizontal and feedback connections. To test this hypothesis, we disentangled the contribution of these signals to anticipatory responses by using the well know intrinsic differences in the spatial and temporal scales of these inputs. To do so, we developed a simple phenomenological, but biologically-plausible, model simulating feed-forward and cortico-cortical responses within a cortical retinotopic map (**Figure 3A**, see Methods and **figure supplement 3**).

**Figure 3.**
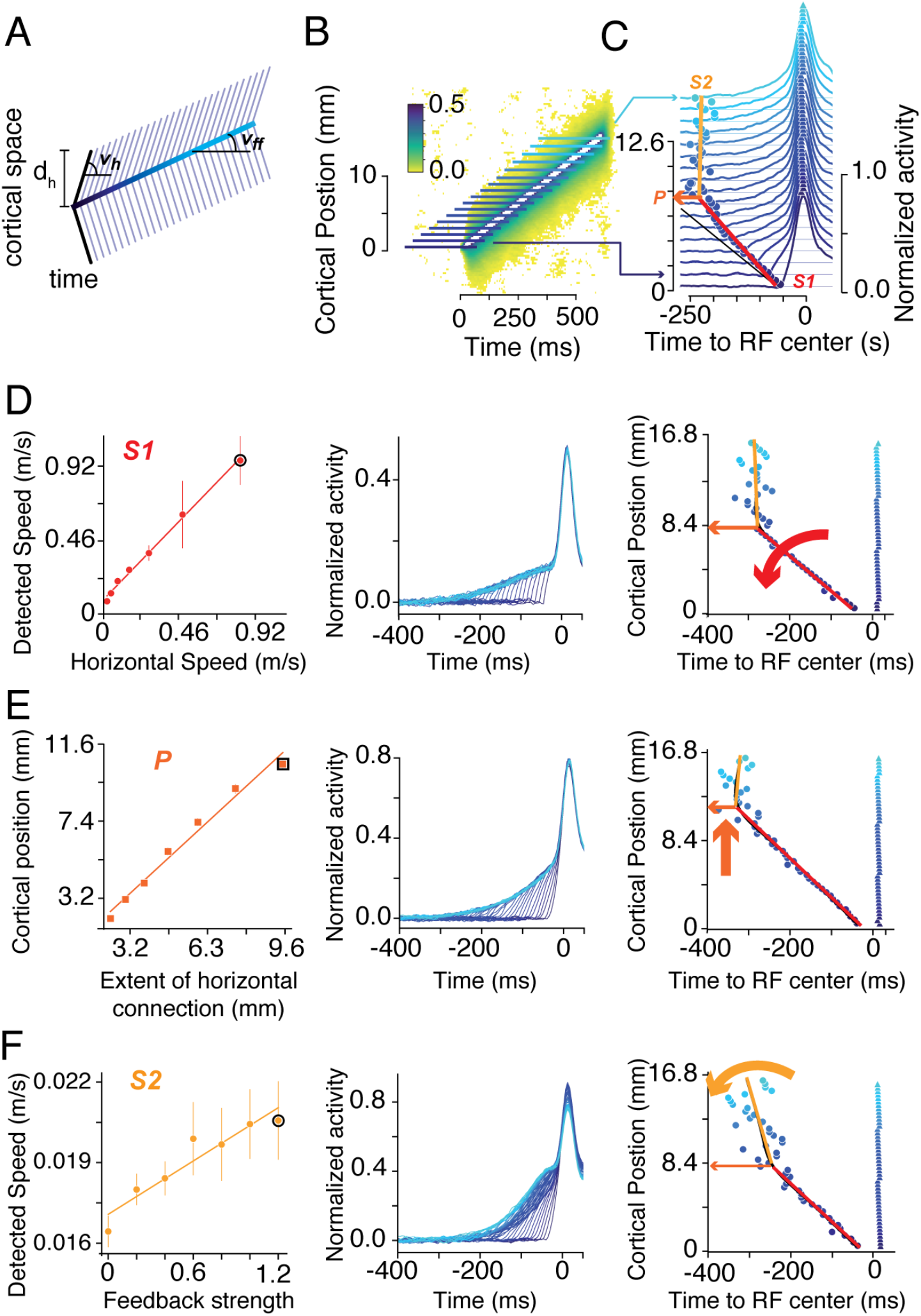
A physiologically plausible model of lateral cortical integration. (**A**) Cartoon describing the model developed to predict the time-course of the neural population response to a translating bar in the space-time domain. The translating bar evokes a sequence of spatio-temporal feedforward activation of the cortex at a speed of v_ff_ (thick blue line, blue hue codes for time). The feedforward activation is then relayed by horizontal propagation at speed *v_h_* and over a distance of *d_h_* represented by thinner obliques lines. (**B**). Activation of the model when *v_ff_*, *v_h_* and *d_h_* are set to values compatible with our visual stimulation paradigm and the know horizontal network properties of the cortex (Angelucci et al., 2002; Girard et al., 2001; Grinvald et al., 1994; Muller et al., 2014). Color-code represents cortical activation strength (arbitrary units). (**C**) Time-course of the neural population response at different positions along the trajectory. Traces are aligned to the time of the feedforward arrival in the receptive field centre (the stimulus onset is displayed as a slant dark line). For each response time-course, latency is indicated by a disk. The profile of latency changes with the increase of the trajectory length is captured by a piecewise linear regression (red and orange lines) showing a first decrease with a slope S1 up to a position P, followed by a second slope S2. (**D**) Effect of changing the speed *vic* on the slope S1 (left). The middle and right panel shows, for a speed example (black circle in the left panel), an example of time-courses and the plot of latencies as a function of distance along the trajectory (with extraction of S1, P and S2). (**E**) Same as in D but for changes of *d_ic_*. (**F**) Same as in D but for changes of feedback strength.

The response to a bar translating along an extended, straight, trajectory was modelled as a sequence of feedforward activations at different spatiotemporal positions, changing with speed ***v_ff_*** (**Figure 3A**). At each spatiotemporal position, this feedforward input also gave rise to an isotropic horizontal propagation of activity with speed ***v_h_*** over a cortical distance ***d_h_*** (**Figure 3A**). When ***v_h_*** > ***v_ff_***, the model qualitatively predicts the emergence of a gradual anticipatory build-up of activity whose onset latency depends on trajectory length (**Figure 3C**). This effect is a consequence of the speed difference between the sequence of feedforward activations (i.e. stimulus speed) and the horizontal propagation: after an initiation phase, a given cortical position will receive information about the approaching bar first from the intra-cortical propagation (i.e. anticipatory response) and second from the feedforward input. For each position, after realigning the time-course of activation patterns to the end of all trajectories, we observed a gradual emergence of an anticipatory activation, whose latency (blue dots) decreases with trajectory length (color-coded as hue of the blue) and reaches saturation for distances larger than *d_h_* (**Figure 3D**). Beyond this distance, the latencies of anticipatory responses do not change anymore simply because of the absence of direct anatomical horizontal connections. In other words, to generate a latency decrease, the bar needs to reach a region that is directly connected to the recording site.

To capture this behavior, we used a piecewise linear regression of the latency as a function of trajectory length. From this regression, we extracted the slope of the change in the response latency at the beginning and at the end of the trajectory (**S1** and **S2** respectively) and the position of intersection between these regression lines (**P**). These 3 parameters correspond to the speed at which latency of anticipatory activity decrease along the trajectory (S1), the spatial limit of this decrease (P) and the rate of latency changes for distances beyond this position (S2). To understand how does the spatio-temporal properties of the intra-cortical propagations affect these three parameters, we varied *vh* and *d_h_* over a large range of values. As predicted, we found that **S1** and **P** are under control of *vh* and *d_h_* respectively, which linearly modulate these parameters over a large range of values (**Figure 3D and E**). In short, the properties of the intra-cortical propagation directly influence the speed and the extent of the latency decrease.

This observation has two theoretical consequences. First, if anticipation would result from a large divergence of the feedforward inputs, we should observe a latency decrease at a fixed delay after stimulus onset since all thalamic input arrive with the same latency. As a result, anticipation should not be trajectory-dependent and should generate a very high speed S1 (**Figure 3D**). Second, the spatial extent of the observed latency decrease should be equivalent to anatomical extent of the anatomical divergence (**Figure 3E**). In the following section, we tested the validity of this model with further neural recordings at different scales.

### VSD imaging unveils an intra-cortical propagation of anticipatory activity

Our model produces testable predictions for the spatiotemporal profile of V1 population responses to a translating stimulus. To test these predictions, we need to record cortical activity at both high resolution and large field of view in order to measure the fine change of response latency as a function of trajectory length. To do so, we used voltage-sensitive dye imaging (**VSDi**), a mesoscopic imaging technique with high spatial and temporal resolution. Since this technique reveals the aggregate membrane potential oscillations of the cortex (Grinvald and Hildesheim, 2004), we can dissect out the effect of cortical propagation with a high sensitivity. In a third, awake and fixating, monkey (WA), we measured population synaptic activity using VSDi (**Figure 4A**) over a large cortical region representing retinotopically −2dva to 0dva of azimuthal eccentricity.

**Figure 4.**
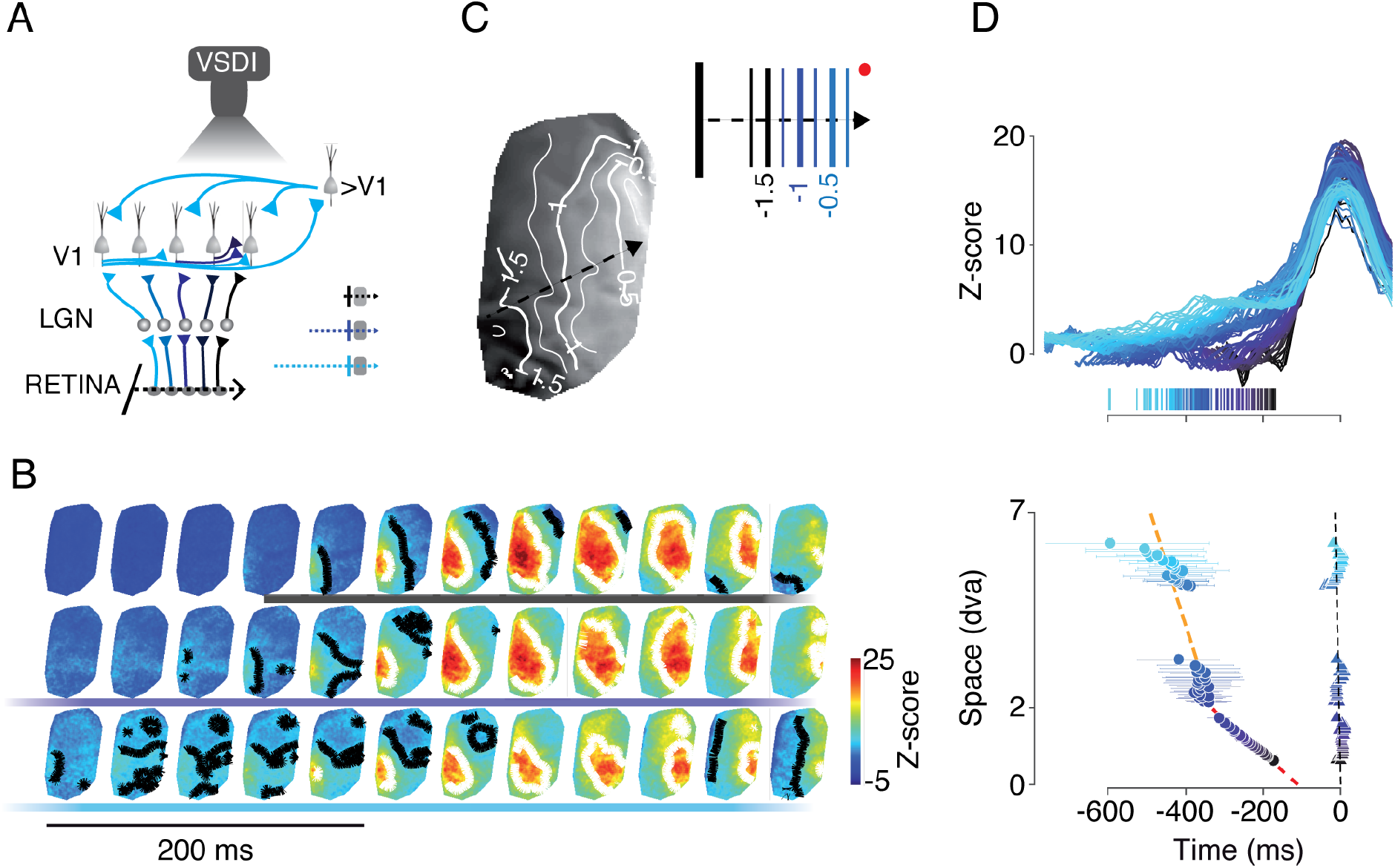
Neural population responses to a translating bar as a function of trajectory length. **(A)** Experimental rationale. Similar to Figure1A but VSD responses are captured using a CCD camera. **(B)** Time sequence of the VSD response map (2 sessions, n=37 trials) for the three trajectory conditions. **(C)** Retinotopic map of the horizontal dimension (grayscale code for the horizontal eccentricity, contours delimit the progression for every 0.5dva). <u>Top-right:</u> Cartoon representing corresponding bar positions in the visual field in the blue color-code. **(D)** <u>Top:</u> Time course of the VSD responses for all conditions broken down by the distance of the initial bar position to the pixel’s retinotopic position. Vertical lines: responses latencies. Color-code same as Figure3C. <u>Bottom:</u> Latency (circle) and time-to-peak (triangle) of VSD responses as a function of the distance of the initial position of the bar to the retinotopic representation (similar to Figure 3C). Red and orange lines are linear regressions.

The translating bar elicited a wave of VSD activity propagating across the retinotopic map at the expected speed and direction of the bar motion trajectory. More important, a strong anticipatory activity was also detectable at the early frames for the medium and long trajectories (**Figure 4B**). Notice that, since the VSDi signal reflects the combination of both the sub and supra-threshold response components (Chemla and Chavane, 2010; Chen et al., 2012), this observation at population level indirectly confirms the spiking results in a third monkey but with a different technique. For similar retinotopic positions, the anticipatory response evoked by the moving bar was stronger for the long trajectory than the one for the medium trajectory (compare first 4 panels in Figure 4B) confirming the slow build-up of activity reported above. Our model demonstrates how such slow dynamics can be a signature of lateral propagation of activity and not a divergent feedforward network.

To facilitate the qualitative comparison with the model simulations shown in **Figure 3C**, we displayed the time course of the VSDi responses as being aligned to the end of the trajectories and sorted accordingly to their distance from the initial bar position (blue hue color code, **Figure 4D**, top panel). For each cortical position, the anticipatory VSD activity started growing slowly a few hundreds of milliseconds before the bar reached the corresponding retinotopic positions. Beyond this point, responses were identical across all conditions, similarly to what was observed at the spiking level, and in close agreement with model predictions. We measured both onset (vertical ticks at the bottom of Figure 4D top panel) and peak latencies for each condition and plotted them as a function of the distance (in visual angle) between the initial position of the bar and the position encoded by each cortical image pixel (**Figure 4D** bottom panel). Responses peaked (triangles) at a fixed delay along the bar trajectory, dynamically mapping the bar motion with the accurate speed (6.8dva/s), corresponding to a cortical speed of 0.016m/s (the equivalent of *v_ff_* in our model). Notice that, in **Figure 4D** we aligned the responses to the end of the trajectories, hereby correcting for bar motion. As a consequence, the vertical alignment of peaks demonstrates that it always occurs when the bar was at the retinotopic position encoded by the retinotopic map (in other words, when it reaches the population RF). In contrast, response onset latencies for pixels close (<2dva) to the starting position of the bar motion decreased rapidly. By fitting this spatio-temporal profile as in the model, we estimated an apparent cortical speed of 0.06m/s, consistent with the speed of intra-cortical horizontal propagation (Bringuier et al., 1999; Girard et al., 2001; Muller et al., 2014; Reynaud et al., 2012). This result suggests that intra-cortical inputs, propagating faster on the cortex than the feedforward activation sequence elicited by the moving bar (0.06 vs 0.016 m/s), is most probably at the origin of the anticipatory activation observed in macaque area V1.

However, for pixels located more than 2dva away from the stimulus starting position (i.e. the position is estimated from the piecewise linear regression fit procedure), responses latencies decreased much less, corresponding to a cortical speed of ~0.02m/s. Such temporal dynamics corresponds to a visual speed of 8.8dva/s, only slightly higher than the bar translation speed (6.6dva/s). Accordingly to our model, such values suggest that the large anticipatory responses seen for long distances, result from the sequential activation of a nested horizontal network driving responses with a spatial range of 2dva. It is compatible with the spatial extent of the horizontal network in macaque V1 (Angelucci et al., 2002). These measured values for both speed and spatial extent are compatible with the intra-cortical connectivity but not with a feedforward origin. Indeed, if anticipation would result from divergence of the feedforward network, our model predicts a faster latency decrease and the spatial extent of this decrease would be smaller than the one we measured (Blasdel and Lund, 1983; Florence et al., 1988; Perkel et al., 1986). Overall, both model prediction and VSDi measurements are in strong agreement to support an intra-cortical origin of anticipatory responses in macaque V1.

What could be the origin of the difference observed between cortical (8.8dva/s) and stimulus (6.6dva/s) speeds? One hypothesis is that it result from gain modulation exerted by long-distance feedback inputs, since only those can mediate such long-range lateral interactions (Angelucci et al., 2002). Therefore, we added a feedback component to our model that acts by varying the relative gain between horizontal and feedforward inputs, as already suggested by others (Liang et al., 2017; Piech et al., 2013; Ramalingam et al., 2013). Increasing the gain of lateral inputs changed the shape of the response time-course (**Figure 3F**) resulting in an increased detectability of the response and, thereby, slightly increasing the slope of the second component in our spatiotemporal profile. These effects are consistent with the results obtained in VSD. To verify this hypothesis, our next step was to test whether the time-scale of such gain modulation was coherent with the dynamics of feedback inputs.

### Low-frequency LFP signals support the existence of an early and diffuse anticipation signal

Convergent studies have recently suggested that the power of low-frequency (i.e. alpha and beta bands) local-field potentials (LFP) signals in V1 can be used as a proxy to quantify the amount of feedback inputs onto V1 (Bastos et al., 2014; van Kerkoerle et al., 2014). Therefore, in order to test our model prediction-that the contribution of feedback signals could participate to the dynamic of anticipatory responses, we studied the time-course of low-frequency LFP signals in V1 in response to our 3-trajectories paradigm (**Figure 5A**). To do so, we measured LFP signals from arrays of 10×10 electrodes (Utah arrays, **MEA**) chronically implanted in two monkeys (BA and PD). The arrays covered a V1 cortical territory of 4×4mm, representing 2-3dva of the motion path. Similar to VSDi, the LFP response (0.3 - 250Hz) to the translating bar across the array followed the projection of the bar to the retinotopic map, swiping across the cortex at the matching cortical speed of 0.015m/s (**Figure 5B**).

**Figure 5.**
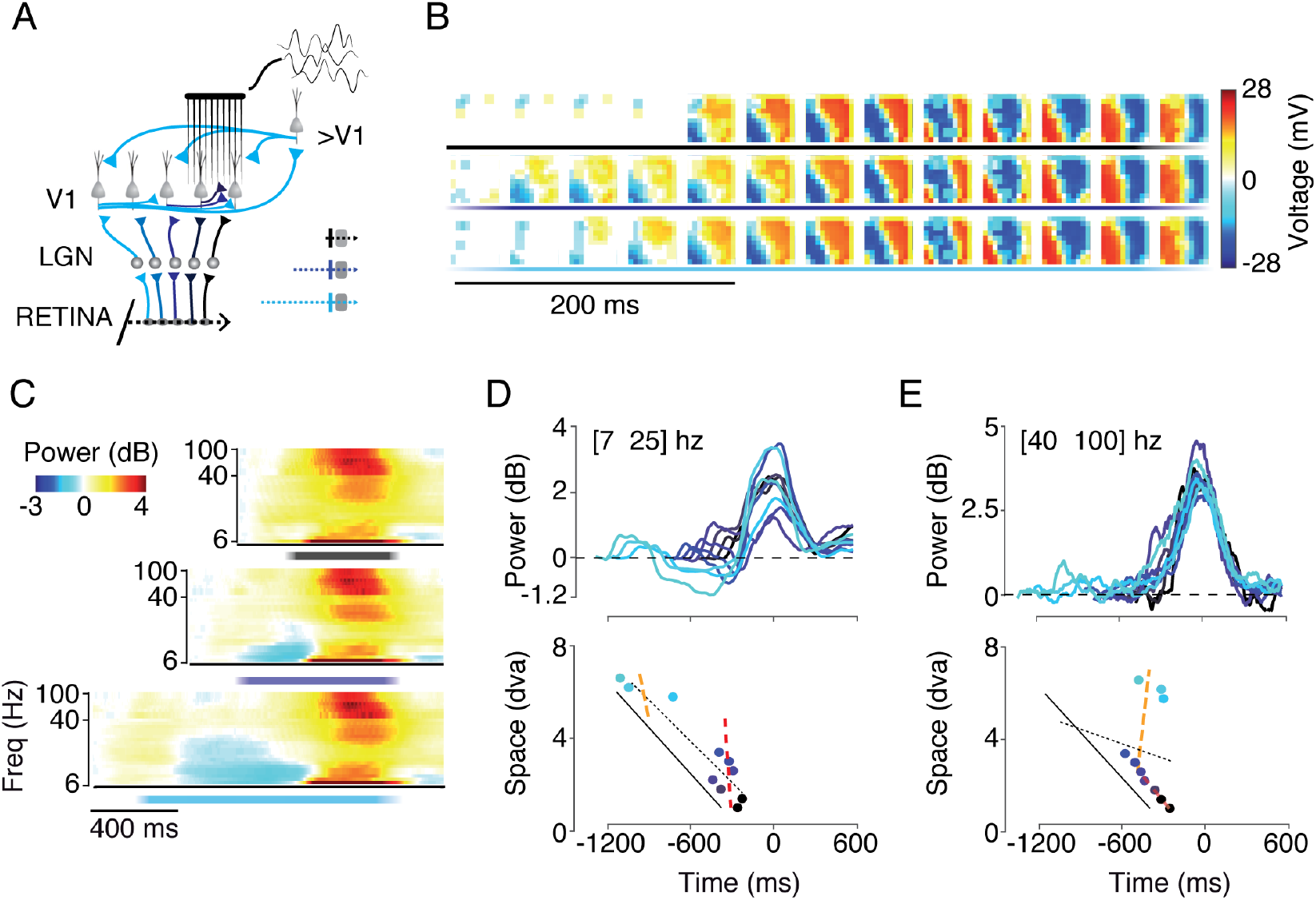
Multi-unit LFP responses as a function of stimulus’ trajectory length. (**A**) Experimental approach. Similar to Fig. 1A but the LFP of the neural population response is recorded using a square multi-electrode-array (MEA) (**B**) Time sequence of the map of LFP responses in MEA space for the three trajectory lengths, (3 sessions, 52 trials per condition). Colored lines at the bottom indicate stimulus presentation and condition type (color-code) (**C**) Time-frequency representation of the LFP power averaged across 96 electrodes. (**D-E**) <u>Top:</u> Time-course of the averaged LFP power for low (D) and high (E) frequency bands, for all conditions broken down by the distance of the initial bar position to each electrode retinotopic position (t=0ms). Black-to-cyan color codes for the increasing distance of the electrode retinotopic position to the initial position of the bar. <u>Bottom:</u> Latency (circle) of LFP responses as a function of the distance of the initial position of the bar to the retinotopic representation. The black line corresponds to the bar trajectory time-course. The orange and red dotted lines are the result of a piecewise linear regression and the black dotted line of a single linear regression.

For longer trajectories (lower rows), clear anticipatory LFP responses were observed all over the array. As for above, we aligned the responses from all electrodes based on their retinotopic position and calculated the time-course of the LFP power-frequency spectrum for the responses to the three trajectories conditions (**Figure 5C**). The anticipatory response was evidenced as a power increase or decrease at high or low frequencies, respectively. Across all frequencies, the response peak always occurred when the bar crossed the centre of the retinotopic area covered by the electrodes array. To detail the dynamics of low and high frequency LFP signals as a function of the bar trajectory length, we filtered the LFP responses using two bandwidths (low: bandwidth 7-25Hz and high: 40-100Hz) and aligned these signals based on the distance between the RF position for each electrode and the initial position of the trajectory (**Figure 5D-E**, top panels). We found a very early and strong anticipatory decrease in the low frequency band. At higher frequencies, however, the anticipatory build-up was more similar to the spiking activities described above, both in terms of amplitude and latency. For both frequency bands, changes were graded according to the length of the motion path. To interpret these results in the framework of our model, we extracted the latency of low- and high-frequency bands and plotted them as a function of trajectory length (**Figure 5D-E**, bottom panels). At high frequencies, in the gamma-band regime, we observed a dynamic similar to single-units and VSDi with an initial phase of latency decrease well captured by a propagation of 0.14m/s. For distances larger than 2dva, the decrease in latency is stable. However, at low-frequencies, in the alpha to beta-band regimes, we observed a constant delay after stimulus onset for all trajectories.

This temporal dynamic could be due to divergent inputs from either thalamo-cortical or feedback pathways, since both can generate synchronous inputs over a large extent. However, given the limits on the spatial extent of the thalamic axonal arbors in primates (less than 0.5mm, (Blasdel and Lund, 1983; Florence et al., 1988; Perkel et al., 1986)) and the divergence of thalamocortical inputs (less than 0.5mm in diameter, (Perkel et al., 1986)), this effect is more probably due to feedback from downstream areas that, in contrast, have the appropriate spatial and temporal scales (Angelucci et al., 2002; Girard et al., 2001; Stettler et al., 2002). Overall, these results are consistent with the predictions of our model. In the last part of our study, we further support the hypothesis that the generation of anticipatory responses necessitates lateral propagation of activity using a complementary approach.

### Anticipation is cancelled for inter-hemifield stimulation

In all conditions tested so far, bar trajectories were all presented in the contra-lateral visual hemi-field, activating a continuous cortical path confined within the recorded hemisphere. If the intra-cortical origin hypothesis holds true, anticipatory responses should not be evoked when the trajectory has its origin within the ipsilateral visual hemifield. This is due to the fact that, since our recordings are retinotopically located very close to the vertical meridian, such stimuli would activate first the opposite hemisphere, generating activation to the recorded location through the myelinated callosal V1-V1 connections, that are much faster than the amyelinated lateral connections and are mostly restricted to a region close to the vertical meridian (Kennedy and Dehay, 1988; Kennedy et al., 1986).

In three monkeys, we tested this prediction using both LFP (**Figure 6A**) and VSDI recordings (**Figure 6B**). As expected, for the opposite stimulus direction, the peaks of the LFP and VSDi responses move towards the opposite cortical direction (compared to **Figures 4** and **5**). However, in stark contrast to a stimulus restricted to the contra-lateral hemi-field (**Figures 4, 5**), moving a visual stimulus from the ipsi-lateral hemifield did not generated anticipatory responses, for both types of recordings and for both low and high frequency power values (see below). This result provides another strong evidence that the observed anticipatory responses are mainly driven by propagations within the intra-an inter-cortical, horizontal and feedback networks.

**Figure 6.**
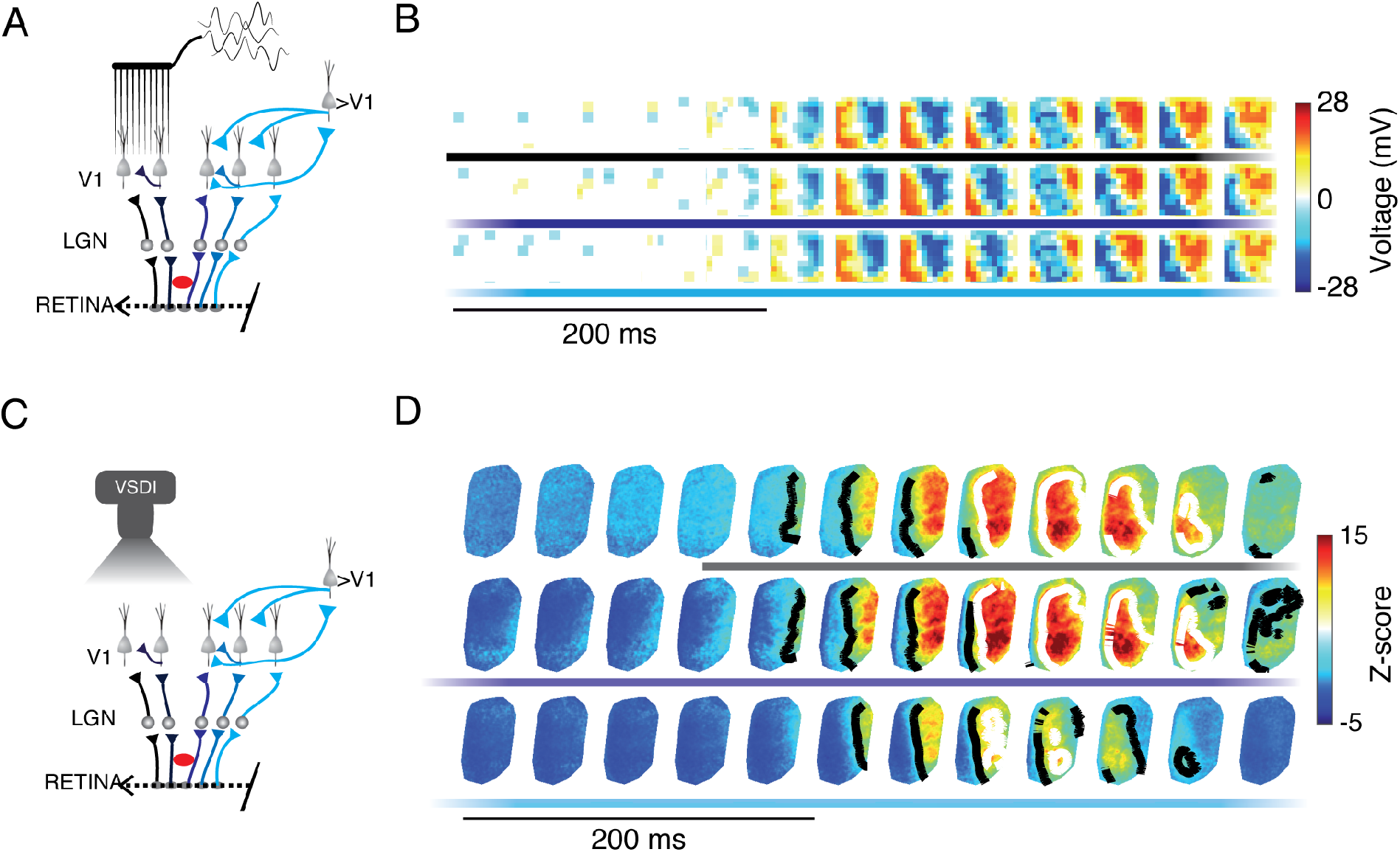
Neural population responses to trajectories with different lengths presented in the ipsilateral visual hemi-field. (**A,C**) The experimental rational for LFP (A) and VSDi (C) recordings. Neural responses are measured in the retinotopic area close to the vertical meridian for a bar starting to move from the ipsi-lateral visual hemifield (i.e. activating the opposite hemisphere) and finishing its trajectory just after crossing the vertical meridian (i.e. activating the recorded hemisphere). Red discs indicate the position of the fixation spot. (**B**) and (**D**) same legend as in Figure 4B and 5B respectively.

Lastly, we compared the average responses latencies across all monkeys, for the three complementary recordings, the three different trajectory length and the two trajectory directions (**Figure 7**). For a stimulation within the contra-lateral hemifield, anticipation gradually develops along the trajectory length in the various measures of neuronal activity. The strongest anticipation was seen for low-frequency band LFP responses (squares), followed by VSDi (triangle) and high-frequency LFP (diamond) responses, leading to a final anticipation expressed at suprathreshold level in single unit activity (circle, **Figure 7A**). For the opposite direction, in both LFP and VSDi recordings, no sign of anticipation is evident for both LFP and VSD signals. Moreover, delayed responses are observed in the VSDi responses for the long trajectories. (**Figure 7B**).

**Figure 7.**
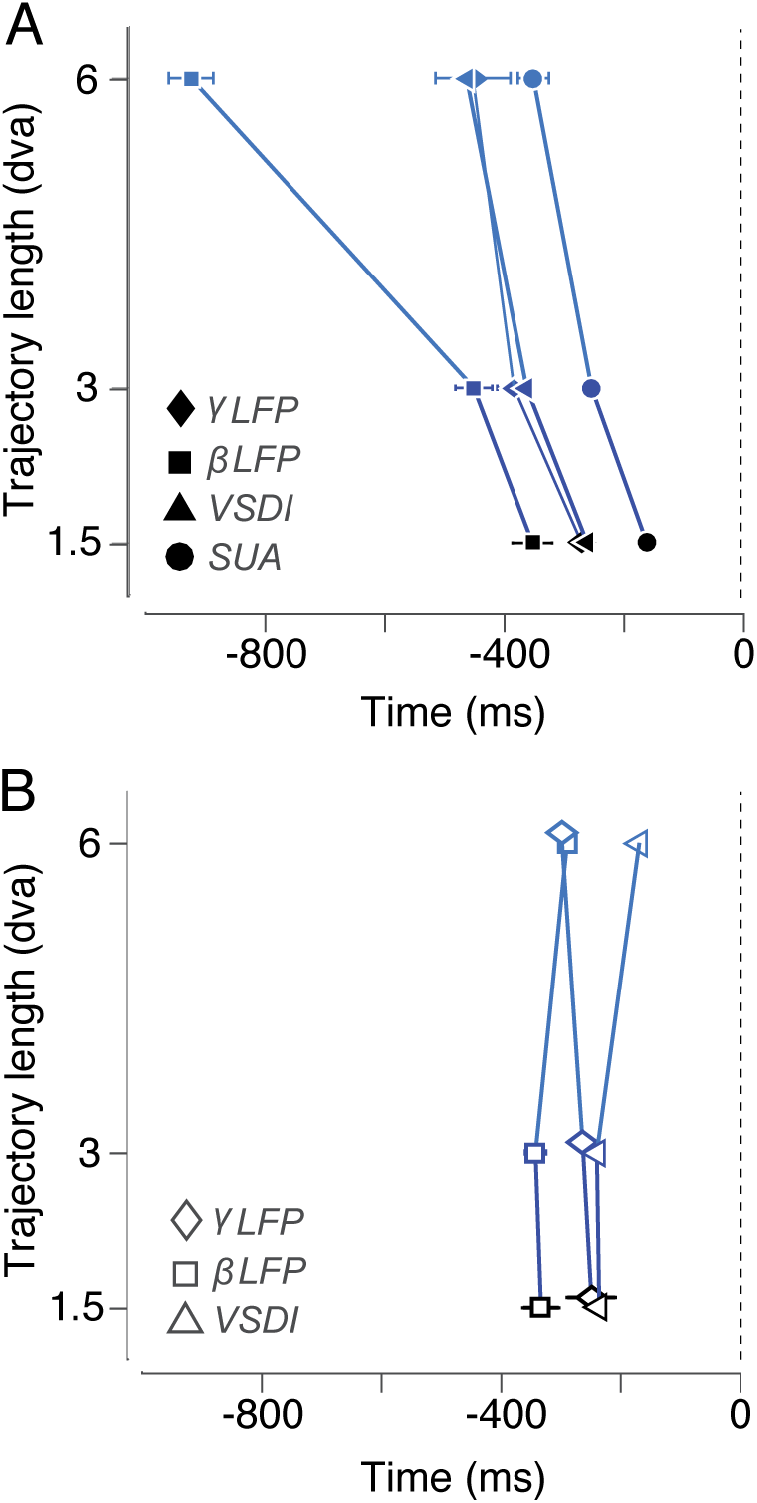
Comparison of results across all recording techniques and 4 monkeys. (**A**) Averaged latencies of SUA (2 monkeys), LFP power and VSDi responses as a function of trajectory length for stimulation within the contra-hemifield. (**B**) Same than A but for a bar coming from the ipsi-lateral hemifield. SEM is calculated across all neurons (SUA) electrodes (LFP) or pixels (VSD).

## DISCUSSION

In primary visual cortex of awake, fixating monkeys, we show that a bar translating along a smooth and extended trajectory significantly activates half of the recorded neurons several hundreds of milliseconds and a few degrees before actually entering their classical RF. This trajectory-dependent anticipatory activity has a specific slow temporal profile composed of a ramping response that is intrinsically different from the classical transient and fast responses driven by pure feed-forward cortical inputs. Such anticipatory activity cannot be explained by an underestimation of the RF sizes or by the scatter in the eye positions. These surprising anticipatory spiking responses present a wide range of timings across neurons, longer trajectory lengths leading to earlier response build-up onsets. This variability in anticipatory response onset is partially accounted for by the neurons’ direction preference: cells preferring the motion direction of the bar display the strongest and earliest anticipatory build-up of spiking activity. In contrast, after entering the classical RF of the neurons, the translating bar produces the same spiking activity regardless of its trajectory length and therefore its cortical mapping history.

Anticipatory responses in the visual system have already been reported in a few physiological studies, first in vitro in the retina of rabbit and salamander (Berry et al., 1999), in area 17 of anesthetized cat (Jancke et al., 2004b; Orban et al., 1985), but also monkeys areas V1 (Guo et al., 2007; Subramaniyan et al., 2015) and V4 (Sundberg et al., 2006). However, these previous observations all differ on several key features when compared to our present study. First, these previous studies (with the exception of Guo et al., 2007, see below) were inspired by the flash-lag effect, in which a translating stimulus is judged ahead of a stimulus flashed at the same position at the same time. In this context, these studies have compared the latency of responses evoked by bars in translation to bars flashed alone in the RF centre. The RF was therefore stimulated differently by the flashed and the moving stimuli, in the sense that the moving stimulus activated a sequence of positions in the RF before reaching the position in which the flashed stimuli was presented for comparison. In contrast, in our approach, the classical RF is stimulated exactly in the same way for all trajectory conditions. Consistent with their initial hypothesis, these previous studies showed a global shift in the time-course of their responses, while maintaining the same shape (i.e. time-to-peak and response offset present similar anticipation than response onset). In our study, anticipatory responses consist in the appearance of an early ramping activity for long trajectory. Afterwards, the response peaks and decrease at the same time than shorter trajectories.

Our results are in accordance with an earlier study carried out in the anesthetized macaque on a small number of neurons, where a bar flashed just outside the classical RF of a V1 neuron can evoke a response when presented after a sequence of 3 other collinear bars in apparent motion, outside the RF (Guo et al., 2007). However, the phenomenon reported in this study is again different from the one we report, since the apparent motion stimulus used will stimulate V1 in a sequence of discrete and transient feedforward activations, creating an intrinsically different activity pattern than that caused by a smooth translating stimulus. Indeed, it has been recently demonstrated, using VSDi in the awake monkey, that a two-strokes apparent motion stimulus generates two waves of propagation interacting non-linearly, instead of many cumulative propagations produced by each position visited by a translating object (Chemla et al., 2019).

Crossing different physiological signatures at single cell and population levels, and guided by a computational model, we provide convergent evidence that the observed spiking anticipation is not a mere consequence of similar events occurring at earlier processing stages but rather results from a specific propagation of activity within V1 intra- and inter-cortical networks. Our biologically realistic-model shows that the cumulative recruitment of converging horizontal input can produce a gradual build-up of anticipatory activity preceding the feedforward activation. Such build-up is evident at both sub-threshold and spiking levels. The trajectory-dependent shift in response onset timing was also observed with high resolution and large field-of-view using VSDi. From the time-course of these neural population responses across the retinotopic map, we have been able to estimate the speed and spatial extent of the propagation underlying the observed anticipation. The spatio-temporal properties of such propagation are fully compatible with the well-known characteristics of the cortical horizontal network in primate area V1 for trajectories up to 2dva (Angelucci et al., 2002; Girard et al., 2001; Muller et al., 2014).

On the other hand, the further decrease in the response onset we observed using VSDi for trajectories extending beyond 2dva (i.e. covering more than 6mm of cortical space) cannot be explained by this horizontal network alone because of the limitation in horizontal connections length. Therefore, we hypothesized that a divergent feedback input from higher visual areas could generate such fast and long-distance effect (Angelucci and Bressloff, 2006; Bair et al., 2003; Bullier, 2001). Using our model, we found that a feedback mechanism can indeed induce a further, though smaller, decrease of response onset for these larger trajectory distances, as observed with VSDi. To support this hypothesis, we show that the bar appearance, even for trajectories longer than 2dva, triggers a fast and strong decrease of low-frequency LFP power, that is likely to indicate a feedback modulation (Bastos et al., 2014; van Kerkoerle et al., 2014). Furthermore, the spatial and temporal properties of this signal are compatible with the known properties of the feedback connectivity (Angelucci and Bressloff, 2006; Bair et al., 2003; Bullier, 2001; Girard et al., 2001). As suggested for sensory-motor networks, such decrease could be related to reporting an expected change in the input, a deviation from the “status quo” (Engel and Fries, 2010).

A last and strong piece of evidence in support of the intra-cortical propagation mechanism comes from our control experiment in which we moved the bar from the ipsilateral visual hemifield, first activating the cortical hemisphere opposite to our recordings. In fact, while slow lateral interactions, mediated by amyelinated horizontal connections, are restricted to the same hemisphere, inter-hemispheric interactions, mediated by fast myelinated callosal connections, are restricted to a region of the visual field close to the vertical meridian (Kennedy and Dehay, 1988; Kennedy et al., 1986). Therefore, a bar moving within the ipsi-lateral visual hemifield would not activate the horizontal network of the recorded cortex. Fast, inter-hemispheric callosal interactions will signal the arrival of the bar in the contra-lateral visual hemifield but only when it approaches the vertical meridian and without any precise retinotopic projection. This is consistent with fact that we did not observe any anticipatory responses in our population activity measurements (VSDi, LFP) for ipsi-lateral approaching bar. While we did not record single unit activities under this condition, VSDi and gamma power LFP signals indicate also the presence of spiking activities. We shall therefore not expect to find any significant slow ramping of spiking activity when a moving object approaches from the other side of the vertical meridian. This does not preclude, however, that inter-hemispheric trajectories could be linked at other spatiotemporal scales in extra-striate cortex that can exhibit larger receptive fields with ipsilateral representations (Desimone et al., 1993; Gross et al., 1977; Pigarev et al., 2001).

All these converging evidences thus suggest that trajectory-dependent, anticipatory, spiking activity results from the interplay of intra- and inter-cortical networks, both propagating activity faster than the feedforward sequence of inputs. Incidentally, our results also show that, through the accumulation of convergent sub-threshold activations, intra-cortical network can drive spiking responses several hundreds of millisecond before the arrival of feedforward inputs, instead of being only modulatory (see also (Jancke et al., 2004a). A possible functional consequence could be that V1 neurons can signal the direction of a moving object before it reaches their receptive fields. Indeed, the fact that anticipatory responses are stronger for neurons whose direction preference matches the trajectory motion direction suggests that direction-selective anticipatory signals can reach downstream populations of neurons (for example MT cells). Thus, motion integration might be facilitated by propagation of anisotropic and anticipatory signals, alleviating motion uncertainties (Burgi et al., 2000; Perrinet and Masson, 2012; Tlapale et al., 2010).

Our study documents for the first time that the neural activity evoked along the trajectory travels across the retinotopic map at the cadence imposed by the FF response, representing at each moment in time information about the stimulus recent history but also its likely future location and velocity. The cortical representation of the leading edge of a motion trajectory is gradually spreading out over a larger retinotopic area than stimuli presented along short trajectories. Downstream extra-striate visual neurons with large RFs could therefore integrate both the feedforward evoked population response with this trajectory-dependent population response. This could be relevant in explaining why stimuli presented at the leading edge of motion trajectories are more detectable (Arnold et al., 2014; Lenkic and Enns, 2013; Liu et al., 2006; Roach et al., 2011; Schwiedrzik et al., 2007) and their location are perceived ahead of a stimulus flashed in the same position (Jancke et al., 2004b; Nijhawan, 1994).

Additionally, this phenomenon could be at the origin of the diffusion of motion information, a mechanism postulated to be necessary for enhancing motion discriminability (Burgi et al., 2000; Grzywacz et al., 1995; Yuille and Grzywacz, 1989) and inferring non-ambiguous motion direction (Anstis and Ramachandran, 1987; Perrinet and Masson, 2012; Ramachandran and Anstis, 1983). Consistently with our observations indeed, such psychophysical effects are modulated by motion trajectory length (McKee and Welch, 1985; Nakayama and Silverman, 1984; Snowden and Braddick, 1989a, 1989b; Watamaniuk et al., 1995). Hence, this circuitry may serve as a plausible mechanism for a predictive coding of future locations and to resolve motion integration computations (Kaplan et al., 2013; Khoei et al., 2017; Perrinet and Masson, 2012). Indeed, such diffusion could subserve two key computational elements of predictive coding: taking advantage of temporal coherency to predict future events and removing highly correlated events within the sensory inflow (Alink et al., 2010; Mumford, 1992; Rao and Ballard, 1999).

In conclusion, our study suggests that propagation of activity within or between cortical areas may play an important role in the dynamical processing of non-stationary stimuli (see also (Chemla et al., 2019; Muller et al., 2018)). We document an effect occurring at the specific spatial and temporal scales of macaque area V1. The anticipatory effect propagates only over a few degrees ahead of the translating bar. We shall envision that the lateral interactions will propagate information across the overlaid functional maps not only in V1, but also in all the other visual areas and maps. In other words, we may imagine a generalization of the effect across the various scales of the encoded retinotopic maps along the visual system, or across different feature maps such as the orientation map in V1 (Chavane et al 2011, Rankin & Chavane 2016, Tlapale et al, 2010), the direction map in MT (DeAngelis and Newsome, 1999) or more complex maps in other extra-striate visual areas. Analyzing systematically the anticipation across these feature maps may help to understand to what extent this anticipatory signal is a simple predictive warning signal or whether it carries more information about the nature of the incoming stimulus. We propose that parallel propagation of information within all these maps, combined with rapid exchange of information between these maps, may be a canonical computational operation by which the visual system process complex, natural, non-stationary inputs.

## METHODS

### 1. Animals and surgery procedures

Experiments were conducted on a total of five hemispheres, in four adult male rhesus monkeys (*Macaca mulatta)*: monkeys NO (right hemisphere), PD (left and right), BM (right), BR (right). Experimental protocols have been approved by the Marseille Ethical Committee for Animal Research (approval #A10/01/13, official national registration #71-French Ministry of Research).

All procedures complied with the French and European regulations for animal research, as well as the guidelines from the Society for Neuroscience. All the monkeys were chronically implanted with a head-holder. Monkey NO, PD and BR were also implanted with a recording chamber located above areas V1/V2. Monkey PD and monkey BM were implanted with a Utah array. Monkey BR had a third surgery for durotomy and transparent artificial dura mater insertion, in order to perform voltage-sensitive dye imaging. Monkeys NO and PD were implanted with a scleral search coil for right eye position monitoring.

#### Multi-electrodes array

Ninety days after head holder implantation, once monkeys PD and BM were trained to achieve a good fixation, a second surgery was performed. Area V1 of the left (monkey PD) and the right hemisphere (monkey BM) were surgically implanted with a 100-electrode Utah array (Blackrock Microsystems, Salt Lake City, UT, USA). The array had an arrangement of 10×10 (PD: Platinum ~400kΩ 1kHz, BM: Iridium oxide ~50kΩ, at 1kHz) electrodes, each of them 1mm long, with an inter-electrode distance of 400μm. The surgery was performed under deep general anesthesia and full aseptic procedures. Anesthesia was induced with 10mg/kg *i.m*. ketamine and maintained with 2-2.5% isoflurane in 40:60 O2-air. To prevent cortical swelling, 2ml/kg of mannitol *i.v*. was slowly injected over a period of 10min. A 2×2 cm craniotomy was performed over the visual cortex and the dura was incised and reflected. The Utah array was inserted in the cortex using a pneumatic inserter (Array Inserter, Blackrock Microsystems) ~5mm under the lunate sulcus, in perifoveal position and covered with a sheet of an artificial non-absorbable dura (Gore-tex). The real dura was sutured back and covered with a piece of an artificial absorbable dura (Seamdura, Codman). The bone flap was put back at its original position and attached to the skull by means of a 4× 40 mm strip of titanium (Bioplate, Codman). The array connector was fixed to the skull on the opposite side with titanium bone screws (Bioplate, Codman). The skin was sutured back over the bone flap and around the connector. The monkey received a full course of antibiotics and analgesic before returning to the home cage.

#### Voltage sensitive dye imaging (VSDi)

The surgical preparation and VSD imaging protocol have been previously described in Reynaud et al., 2012 (Reynaud et al., 2011). Briefly, after head holder and chamber implantation on the right hemisphere, a second surgery was performed, once the monkey BR was trained to achieve a good fixation. The dura was removed surgically over a surface corresponding to the recording aperture (18mm diameter), and a silicon-made artificial dura was inserted under aseptic conditions (Arieli et al., 2002). Such silicon-dura is necessary to have a good optical access to the cortex.

### 2. Recordings

#### Eye Movements

For monkeys NO and PD, horizontal and vertical positions of the contralateral eye to the recorded hemisphere, were recorded using the scleral search coil technique (Judge et al., 1980; Robinson, 1963). For those monkeys, a second surgery was performed to insert a search coil below the ocular sclera, to record eye movements with the electromagnetic technique (Robinson, 1963). For monkey PD and BM we used the Eyelink 1000 (SR Research sampling frequency of 1,000Hz). For monkey BR, we used an ISCAN eye tracking system (512 x 256 x 60Hz, ETL-200).

#### Single electrode (SE)

A computer-controlled microdrive (MT-EPS, AlphaOmega, Israel) was used to insert through the dura a tungsten micro-electrode (FHC, 0.5-1.2MΩ at 1,000Hz) on the right hemisphere of monkey PD and NO. Spikes were sorted on-line using a template-matching algorithm (MSD, AlphaOmega).

#### Multi-electrodes array (MEA)

Recordings were carried out through a 128-channel Cerebus acquisition system. The signal from each active electrode (96 out of the 100 electrodes were connected) was pre-processed by a head stage with unity gain and then amplified with a gain of 5000. The signal was filtered in two different frequency bands to split it into LFPs (0.3–250Hz) and spiking activity (0.5–7.5kHz). The LFPs were sampled at 1kHz and saved on disk. To improve the electrode independence among LFPs, we subtracted from all electrodes the common LFP averaged across all electrodes.

#### Voltage sensitive dye imaging (VSDi)

After removing of the artificial dura-mater, the cortex was stained with the voltage-sensitive dye RH-1691 (Optical Imaging), prepared in artificial cerebrospinal fluid (aCSF) at a concentration of 0.2mg/ml, and filtered through a 0.2μm filter. After a 3 hours staining period, the chamber was rinsed thoroughly with filtered aCSF, to wash off any supernatant dye. The artificial dura was inserted back and the chamber closed with transparent agar and cover glass. Optical signals were recorded from a focal place ~300μm below the surface using a Dalstar camera (512×k512pixels resolution, frame rate of 110Hz) driven by the Imager 3001 system (Optical Imaging). Excitation light was provided by a 150W halogen lamp filtered at 630nm and fluorescent signals were high-pass filtered at 665nm during 1363ms (trial duration).

### 3. Experimental protocols

#### Behavioral task and Training

Monkeys were trained to fixate a red target presented on the center of the screen for all the duration of the trial (~1.5sec) within a window of 1-2dva of diameter. We monitored the eye positions and controlled online the behaviour (sampling rate: 1 KHz) using the REX package under the QNX operating system (Hays et al., 1982). For VSDI experiments, both online behavioral control and image acquisition were heartbeat triggered; heartbeat was detected with a pulse oximeter (Nonin 8600V). After a successful trial, i.e. the monkey has maintained his fixation during the whole acquisition period (~2s), monkeys received a drop of water (NO, PD, BM) or mashed apple (BR) as a reward. For VSDi recordings, an intertrial interval of 8 sec was set for dye bleaching prevention.

#### Visual stimulation

Visual stimulation protocols were displayed using different devices depending on the technique. For both SE and MEA recordings, they have been produced using in-house software (Gérard Sadoc, Acquis1-Elphy, Biologic CNRS-UNIC/ANVAR) but with different displays. In each system, the display was gamma calibrated by means of a lookup table. For the SE experiments, stimuli were back-projected on a translucent screen covering of visual field of the monkey, at a distance of 100 cm, using a video projector (resolution: 1280×1024 pixels at 60 Hz, operating range 0-24 cd/m^2^). The luminance of the motion stimulus was 22.2 cd/m^2^ (background 2.2cd/m^2^). For the MEA experiments, stimuli were presented on a LCD screen 27’ (resolution: 1920×1080 pixels at 60Hz, operating range 0-294cd/m^2^) at a distance of 57cm. Mean luminance of the motion stimulus was 80cd/m2 (background 5cd/m^2^). For VSDI, the visual stimuli were computed on-line using VSG2/5 (VSL v8) libraries on Matlab (The MatWorks Inc., Natick, MA, USA) and were displayed on a 22’ CRT monitor at a resolution of 1024×768 pixels and a refresh rate of 100Hz, operating range 0-75cd/m^2^. The viewing distance was set to 57cm, the mean luminance of the motion stimulus was set to 71cd/m2 and the background was kept constant to 12cd/m^2^

#### SE stimulation protocol

Each neuron was characterized using three different stimulation paradigms. First a sparse noise (SN) paradigm, to quantitatively map out the receptive field (RF) of the neuron. The SN consisted in the sequential presentation of small dark and light squares of 0.6×0.6dva, flashed for 50ms randomly positioned within a 10×10 grid (11.2+/-11 cd/m2, ~15trials, see (Bringuier et al., 1999). The stimulation for the following paradigms was centered on the RF centre (RFc). To estimate the RFc, we extracted the center from the ellipse that was best adjusted to the activity contour at a 3.8 z-score (equivalent to a p-value of 0.1% corrected for 100 positions). Then, a direction tuning (DT) paradigm, using a bright bar (0.5 x 4 dva) moving at 6.6 dva/s. The bar was crossing the stimulation centre in the middle of its trajectory (6 dva altogether) and randomly presented at 12 different directions (spaced by 30° of circular angle). The bar orientation was always perpendicular to the motion direction. Finally, a trajectory length paradigm (3T) in which the same bar, oriented perpendicularly to the motion direction was moved horizontally at 6.6 dva/s, starting at 1.5, 3 or 6 dva from the RF centre and disappearing 1.5 (NO) or 2 dva (PD) beyond the RF centre. The bar trajectory started from a lateral position in the contra-lateral visual hemi-field towards the vertical meridian. In those three protocols, control trials with no stimulation were presented within each block (blank condition). All stimuli were presented in random order.

#### MEA stimulation protocol

Electrodes LFP RF were mapped by the same sparse-noise paradigm used for SE and stimulation was centered on the centre of mass of the population RFs. Then, we used the same protocols (DT and 3T) than for single units recordings. However, we also extended the protocol 3T using a symmetrical trajectory moving towards the RFs from the ipsilateral visual hemi-field.

#### VSDi stimulation protocol

The VSDi stimulation protocol for the 3T paradigm was established using the exact same stimuli (see above). The trajectory vertical position was centered on 2dva below the horizontal, the imaging area being roughly centered on [-2; −1dva] (see retinotopic map using intrinsic optical imaging Figure 4C), while horizontally moving at 6.6dva/s from 7, 4 or 2.5dva on the left of the vertical meridian to 0dva (vertical meridian position). Here again, we extended the protocol using a symmetrical trajectory moving towards the RFs from the ipsilateral visual hemi-field. In addition to these three evoked conditions, two blank conditions, i.e. a grey screen is presented during the acquisition period, were used and all stimuli were randomly interleaved.

### 4. Data Analysis

All data have been analyzed using custom-written routines in Matlab.

#### SUA PSTH plots

For every neuron, we calculated the peristimulus time histograms (PSTH) for the three trajectory lengths conditions, averaging across the trials PSTHs (total number of trials between 15 and 35, bin size of 10ms), smoothed by a low pass filter (window of 120 ms) and expressed in spikes per second. Standard error of the mean (SEM) was calculated across trials. First, the histogram of every trial (bin size 10ms) was smoothed with a low pass filter (window of 120ms). Then we calculated the SEM across these smoothed histograms. Response latency was calculated on PSTHs that had a peak response higher than 1.96 times the standard deviation over the mean of the amplitude of the blank condition PSTH (amplitude threshold). The latency was computed on the PSTH derivative (estimated on each time frame “t” by calculating the linear regression of the PSTH response over 20 points, corresponding to 200ms, centred on t). The latency was defined on each condition as the moment the PSTH derivative was crossing a threshold, (calculated as the mean plus 1.96 times the standard deviation of the derivative of the blank condition PSTH) for 5 consecutive points.

#### Averaged PSTHs

For each monkey, we averaged the PSTHs of cells presenting a statistically significant anticipatory response (two-sample t-test between the distribution of spikes numbers for trial of the longer trajectory length conditions and the short one in a 100ms time window placed just before the response latency for the short trajectory, i.e. just outside the classical RF). Before averaging, single cells PSTHs were aligned on the latency of the short trajectory condition. Second, to remove the unselective component of the response for each cell, the average across time of the PSTH baseline (the first 400ms before stimulus presentation) was subtracted from the full PSTH. To align the PSTHs of each cell on the latency for the short trajectory, for each condition, we first calculated the error between the expected (40ms after stimulus presentation) and measured latency and then we adjusted the PSTHs offset in time to remove it. The statistical significance of the averaged responses was calculated over time (within a sliding time window of 100ms, shifted of steps of 10ms). For each neuron and each condition, the distribution of spikes counts across trials, in each time window, was compared by a two-sample t-test (P<0.05) to the distribution of spikes counts across trials, over all the time windows in the blank condition.

#### Time course of the neurons proportion with statistically significant response

For each neuron, we calculated at which moment in time (within contiguous time windows of 100ms) the distribution of spikes counts across trial, for the evoked conditions, was significantly different from the distribution of spikes counts across trials and all the time windows of the blank condition (one-tailed two-sample t-test, P<0.05). For each time window, we calculated the percentage of neurons with a significant response, with respect to the number of neurons presenting a significant response in the time window corresponding to the RF center (49 neurons).

#### Relative firing rate within a window of 100ms just before and just after the RF border

For each neuron, we calculated the difference in spike rate between the short and the longer trajectories conditions (aligned on the RF centre) within a 100ms window, placed just before the response latency if the response to the short trajectory or just after. The statistically significant response differences between the distribution of spikes numbers for trial of the longer trajectory length conditions and the short one, were calculated by one-tailed two-sample t-test (P<0.05).

#### Neurons preferred direction and direction index

Neurons preferred direction was calculated fitting the neurons tuning curves (built from the PSTH responses within a time window of 200ms centred on the RF centre for all the 12 conditions of the DT paradigm) with a double multiplicative Von Mises function, see (Swindale, 1998). Neuron direction index was calculated as 1, minus the ratio of the response in the opposite direction to the response in the preferred direction.

#### Probability of anticipation

Probability of anticipation was calculated for each cell by one-tailed two-sample t-test (P<0.05) between the short and the longer trajectory conditions (aligned on the latency for the short trajectory condition) within a 100ms window place just before the response latency for the short trajectory.

#### Spikes representation in visual space corrected by eye position deviation from the fixation point (control)

For 22 cells in monkey PD, we checked whether the response anticipation could be explained by eye movements towards the approaching bar. For that purpose, for each trials the time occurrence of each spikes was converted into a visual space coordinate corresponding to the stimulus position in visual space for that particular time assuming a latency of 50ms ((skp timing-50) *stimulus speed). This position was then converted by a position in retinal space taking into account the distance of eye position (low pass filtered across 100 ms) from the fixation centre (the average across all the eye positions), at −50ms from the spike time. Then we calculated the PSTHs for the evoked conditions, in space coordinates, for the rasters before and after correction of spike positions.

#### LFP power spectral analysis

The LFP spectrum (sampling rate 1000Hz) was computed using Chronux open source MATLAB toolbox (http://www.chronux.org/, multi-taper method, *mtspecgramc.m* function). Taper parameters were bandwidth (W): 2.5, taper duration (T): 1, integer (p) such as 2TW-p tapers are used: 2. In order to optimize the frequency/time resolution, we used two different sliding time windows for low (7-25Hz) and high (40-100Hz) frequencies (respectively 400ms and 150ms). The LFP spectrum of the evoked conditions was divided for the mean spectrum of the blank condition (across time) and the response offset at zero frame was subtracted. LFP was recorded through a Utah array in 3 sessions for a total of 52 trials for condition. The spectrogram was obtained by averaging across 96 electrodes, re-aligned on the LFP RF centre of every electrode, determined through the sparse-noise paradigm (as described above).

#### VSD data analysis

Stacks of images were stored on hard-drives for off-line analysis. The analysis was carried out using the Optimization, Statistics, and Signal Processing Toolboxes. The evoked response to each stimulus was computed in three successive steps. First, the recorded value at each pixel was divided by the average value before stimulus onset (frames 0 division), to remove slow stimulus-independent fluctuations in illumination and background fluorescence levels. Second, this value was subsequently subtracted by the value obtained for the blank condition (blank subtraction) to eliminate most of the noise due to heartbeat and respiration (Shoham et al., 1999). Finally, a linear detrending of the time series was applied to remove residual slow drifts induced by dye bleaching (Chen et al., 2008; Meirovithz et al., 2010). For this study, VSD data from two sessions were accumulated by registering them onto a reference session (using a quadratic registration algorithm described by Takerkart et al., 2008(Takerkart et al., 2008)). Then responses amplitudes were normalized to a reference condition and data were *Z*-scored(Jancke et al., 2004a) for further analysis.

#### Retinotopic maps

To obtain a retinotopic representation, we presented the bar stimulus (0.5×4dva) at 6 different static positions (from −2.5 to 0.5dva compared to the vertical meridian) for 600ms (after a delay period of 200ms) and we recorded intrinsic optical imaging signals from the cortical surface(Grinvald et al., 1994). Each trial consisted of 4181ms (92frames) and was followed by an inter-trials interval of 5s. For intrinsic-signal imaging, the cortex was illuminated with a 605nm light. Off-line analysis of intrinsic maps started with the same normalization procedures than for VSD images, i.e. frames 0 division and blank subtraction) providing the cortical activation profile for each static positions of the bar. Then for each pixel, we extracted the mean values in each of the six cortical profiles and computed the centre of mass. The retinotopic map was finally constructed after conversion into visual field positions.

#### VSDI Latency estimations

Response latency was defined as the point in time at which the signal derivative crossed a threshold set at 2.57 times (99% confidence) the SD of its baseline computed during a 100-ms-long window right before stimulus onset.

#### Spatio-temporal phenomenological cortical Model

The model was built to represent V1 activity over a 2D matrix representing retinotopic cortical space in one dimension (full extent of 21mm) and time in the other dimension (full extent of 1sec). In this spatio-temporal representation, we modelled the feedforward input as an activation moving across the matrix at a speed of ***v_ff_*** set to 0.016 m/s (equivalent to 6.6dva/s in cortical coordinate at the eccentricity of our neural recordings). At each moment, the dot-like feedforward input generated an activation profile implemented as a gaussian function in the spatial domain, with a standard deviation of 0.42mm.

For each of these feedforward activations, we computed V1 activity as the interaction of this activation with that of similarly defined horizontal and feedback maps. First, horizontal activity is the result of a spatio-temporal convolution with a kernel (***k_h_***) which is then scaled with a nonlinearity. The kernel is modelled as a spreading gaussian (see eq 1) for which the standard deviation increases linearly with time with a celerity ***c*** and the amplitude decreases exponentially with the square of time relative to ***τ***. The weight ***W_h_*** of this kernel is adjusted to account for the strength of the horizontal input. The speed of the horizontal spread ***V_h_*** was therefore controlled by ***c*** (set by default to 0.1m/s) and the spatial extent is controlled by a trade-off between ***τ*** and ***c***, i.e. the maximum extent being reached for the latest point in time for which the horizontal activity amplitude is still above baseline. As a consequence, we used the value of ***σ*** at time ***t*** =1.5 **τ** (the amplitude will have reach 10% of its maximum) as a proxy for the horizontal extent ***d_h_*** (set by default to 6mm)

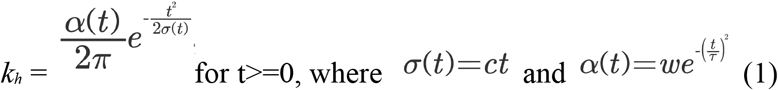

To account for the observation that the feedforward response amplitude does not change along the trajectory, the horizontal activity contribution was scaled through a sigmoid non-linearity (***S***) controlled by the difference between feedforward and horizontal activities. Horizontal and feedforward signals were then linearly combined with a factor of **ρ**.

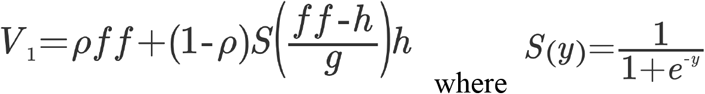

Feedback effect was thereafter modelled as a spatio-temporal multiplicative gain to the horizontal activity. The kernel of the feedback was similarly designed as the one for the horizontal, albeit with much faster speed and larger spatial extent. The resulting activity of the feedback amplified multiplicatively the activity of the horizontal activity (Liang et al., 2017; Piech et al., 2013; Ramalingam et al., 2013). The feedback strength was controlled by a scalar weight (*w_fb_*) that we modulated between 0 (no feedback) and 1.2 (feedback stronger than horizontal).

The evolution of the expected response time-course for different positions along the trajectory can be simply read-out from the matrix representing the spatio-temporal activity. Similarly to what was done in experimental data, latency was calculated as threshold crossing of derivative above a given value (set to 0.01).

## AUTHOR CONTRIBUTIONS

GB and AB performed single unit recordings. GB performed MEA recording. SC performed VSDI recording. FC and LP did the computational model. FC supervised all the experiments and designed the study with the help of GB. GB, GSM & FC wrote the manuscript with the help of AB and SC.

## ACKNOWLEDGMENTS

This work was supported by the Marie Curie Program (FACETS-ITN) the FET Program (FACETS, BrainScales) of the European Union and the ANR Trajectory and Horizontal-V1. We thank Ivan Balansard, Xavier DeGiovanni, Marc Martin and Fréderic Barthélemy for their excellent technical support. We are grateful to Cyril Monier who provided an important help in setting up the MEA experiments and to Lyle Muller for careful reading of the manuscript.

## SUPPLEMENTARY MATERIAL

**Suppl. Figure 1:**
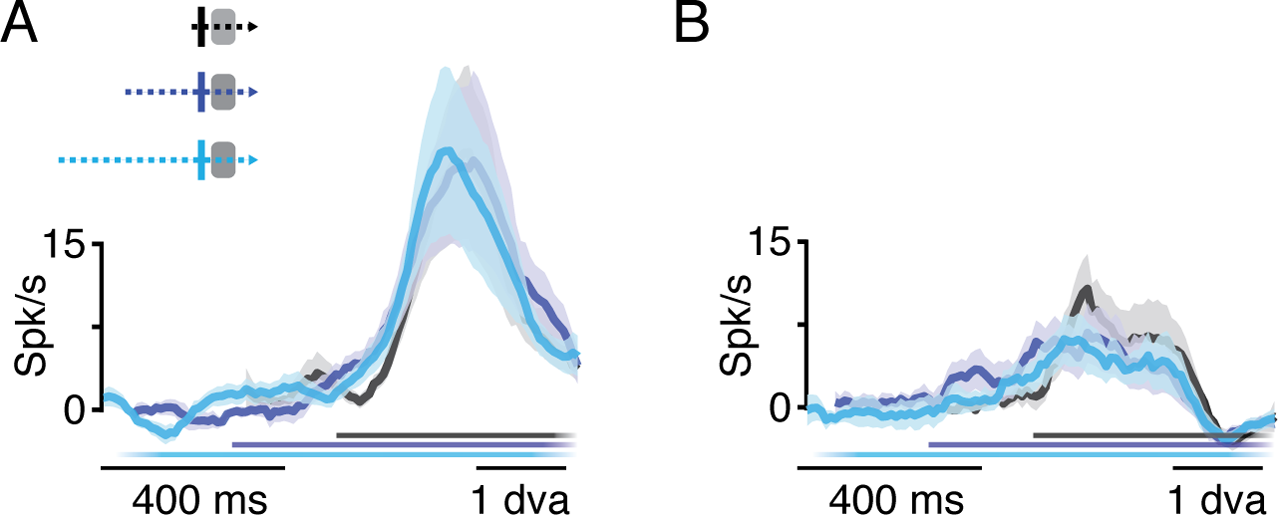
Average PSTHs across non-anticipatory neurons for the three trajectories. **(A)** Average PSTHs across non-anticipatory neurons for the three trajectories for monkey PD (n=20). Responses are aligned to the latency of the short trajectory condition. Color lines (bottom) indicate the stimulus presentation time, for each condition. No significant difference with respect to the short trajectory condition were detected (Two-sample t-test P>0.05). **(B)** Same than A but for monkey NO (n=3).

### Figure supplement 2: Anticipation is not caused by eye movements or excessively large RF

In order to completely exclude potential effects of eye movements or changes in receptive field size to the observed anticipation, we conducted several control experiments. First, we computed the distribution of eye positions across all successful trials of the 3-trajectories paradigm for 22 cells (a smaller number of cells is used here because the eye-movements recorded in some experimental sessions were deteriorated due to an error in the data acquisition software). Our results show that 99% of eye positions were distributed within 1dva around the fixation spot and with identical distribution for the 3 conditions (**Supp. Figure 2A**). Thus, any changes in RF location due to eye movements would have been much smaller than the 2.3 (PD) and 3 dva (NO) equivalent distance reported for the average anticipatory responses in the two monkeys.

Second, comparing the skewness of the distributions in Figure 1A for long (y-axis) and short trajectory conditions (x-axis), we showed that there is not difference in the slant towards of (negative values) or away from (positive values) the direction of the approaching bar (**Supp. Figure 2B**).

Third, we recomputed mean response profiles as a function of the actual distance of the bar from the RF center when considering the real-time position of eye fixation (acquisition rate 2Hz) (**Supp. Figure 2C**). This analysis (A-C) was applied to the 22 cells for which we had high quality eye-movements recordings. The anticipatory responses for medium (3dva) and long (6dva) trajectories were still significant in all the anticipatory cells tested (n=7), rejecting the possibility that they could be a mere consequence of a systematic bias or transient events in fixation behavior. On the contrary, the overall response across the 22 cells tested presents an increase in the anticipatory response after correcting for the real-time position of the eye.

Fourth, we confirmed that anticipatory responses are not simply explained by a sampling bias for large receptive fields (Supp. Figure 2D, E, F). The anticipation is plotted in spatial coordinate as a function of the receptive field radius for medium (1D) and long (1E) trajectories. The anticipation started systematically outside of the classical RF (above the main diagonal) and most of the points are also further away than twice the size of the classical RF (above y=2x). We also compared the p-value of the statistical test for the significance of the anticipatory response (relative to baseline) and showed that there is not trends, actually the smallest p-values are observed for the smallest RF.

**Suppl. Figure 2:**
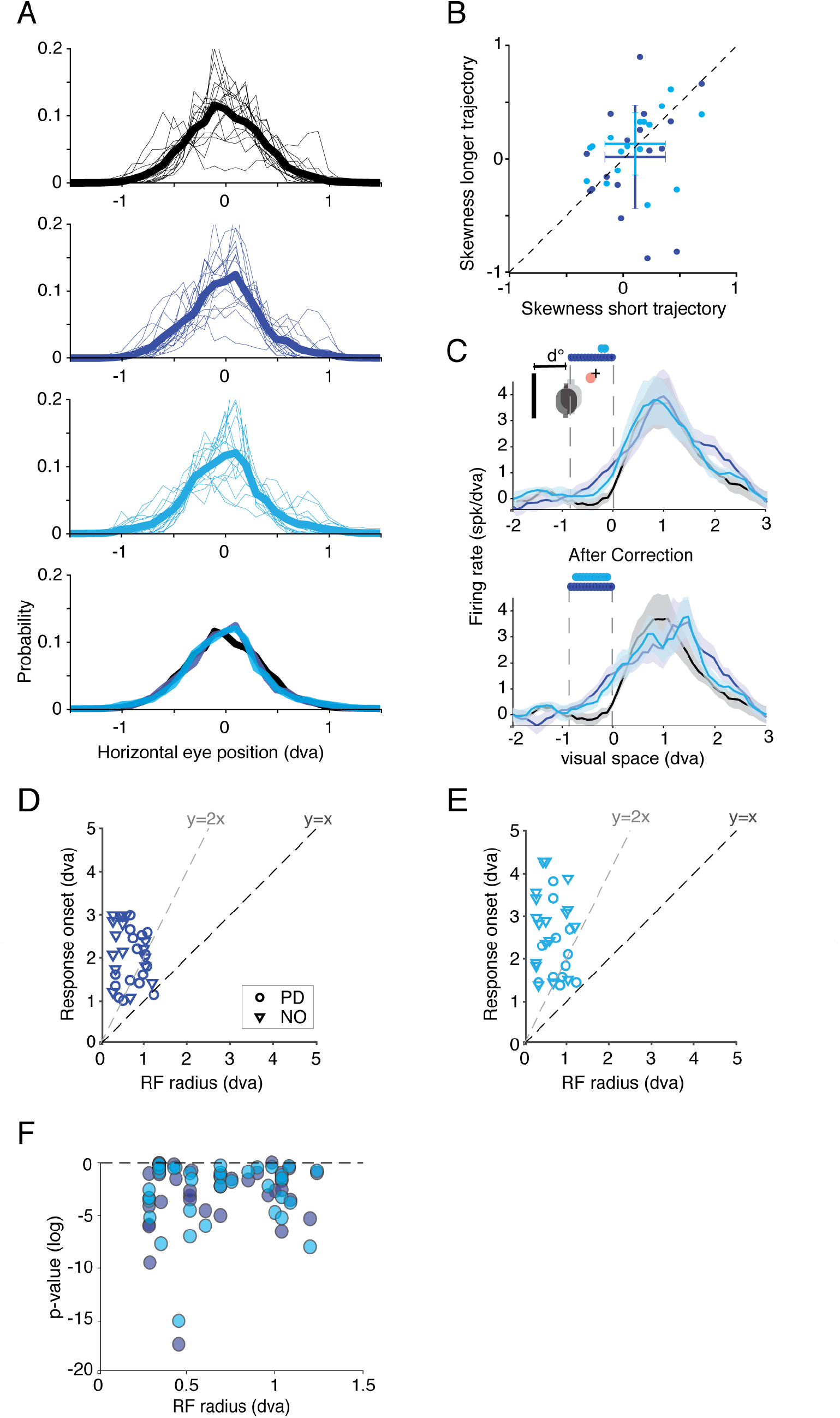
Test existence of biases from eye movements and RF sizes in the emergence of anticipatory responses. **A**) Gaze positions distribution across all successful trials for all 3-trajectories conditions (black), thin traces are single cells, thick trace the average, which are plotted for the 3 conditions in the same graph in the bottom (n = 22 cells). (**B**) for all cells we measured the skewness of the distribution in A to test whether a slant towards (negative) or away (positive values) of the direction of the approaching bar was existing for long trajectories (y-axis) compared to the short trajectory condition (x-axis). The average and std across cells is presented as error bars. (**C**) Eye movement correction control. Average PSTHs among cells (n=22 cells) for the three conditions before and after eye movement correction of the spikes timing relative to the exact distance between the receptive field and the bar. Neural response is therefore expressed in number of spikes per degree of visual angle of the bar displacement (spk/dva), assuming a constant latency of 50ms. Upper left inset sketch the RF position correction (grey before, dark after correction) due to eye position (red dot) relative to the central fixation (‘+’), allow to compute the exact distance (d in dva) between the bar and the RF. Colored lines (top) indicate significant difference with respect to the short trajectory condition (two-sample t-test P<0.05) before the bar enters the RF (i.e. 0 dva). (**D-E**) Position of the bar at response onset (y-axis) as a function of RF size for the medium and long-trajectory conditions. The main diagonal (y=x) and twice the main diagonal (y=2x) are shown to compare the position of the bar that evoked anticipatory activity with extreme limits of RF when mapped using classical procedures. (F) p-value of the anticipatory activity t-test (relative to spontaneous activity) as a function of its RF size (n = 36).

**Suppl. Fig 3:**
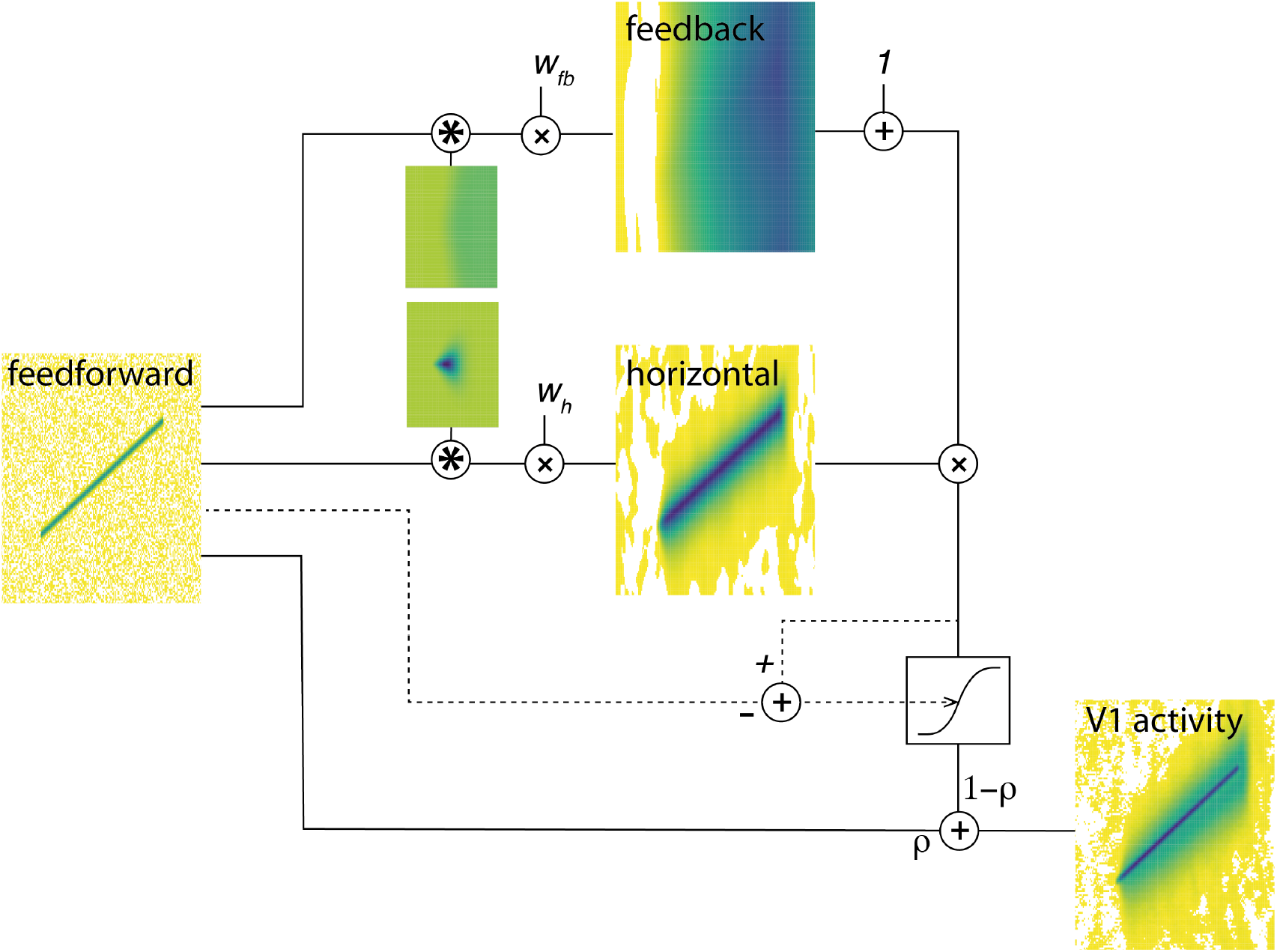
Block diagram sketch of the computational model. From left to right is depicted the logical flow of our model. The matrices of the diagram all represents space in y and time in x. On the left is represented the feedforward predicted activation of V1 to a bar moving at 6.6dva/s other an extended trajectory. At that level, random noise is added to the simulated activation. To generate horizontal or feedback activation maps, the feedforward matrix is convolved with kernels with low extent low speed (horizontal) and large extent fast speed (feedback), and then weighted by *w_h_* and *w_fb_*. The feedback activation map is then added a scalar of 1 and multiplied to the horizontal activation map to amplify of the horizontal activation (Liang et al., 2017; Ramalingam et al., 2013, Piech et al 2013). The resulting activation map is then passing through a sigmoid non-linearity controlled by the difference between the feedforward and the horizontal-feedback activation maps. The relative contribution of the feedforward and horizontal-feedback maps are then summed with a ratio of p, to generate the predicted V1 activity.

